# Structural basis of transcription-translation coupling

**DOI:** 10.1101/2020.03.01.972380

**Authors:** Chengyuan Wang, Vadim Molodtsov, Emre Firlar, Jason T. Kaelber, Gregor Blaha, Min Su, Richard H. Ebright

**Affiliations:** Waksman Institute and Department of Chemistry and Chemical Biology, Rutgers University, Piscataway NJ 08854, USA; Rutgers New Jersey CryoEM/CryoET Core Facility and Institute for Quantitative Biomedicine, Rutgers University, Piscataway NJ 08854, USA; Departent of Biochemistry, University of California, Riverside CA 92521, USA; Life Sciences Institute, University of Michigan, Ann Arbor MI, 48109, USA

## Abstract

In bacteria, transcription and translation are coupled processes, in which movement of RNA polymerase (RNAP) synthesizing mRNA is coordinated with movement of the first ribosome translating mRNA. Coupling is modulated by the transcription factors NusG--which is thought to bridge RNAP and ribosome--and NusA. Here, we report cryo-EM structures of *Escherichia coli* transcription-translation complexes (TTCs) containing different-length mRNA spacers between RNAP and the ribosome active-center P-site. Structures of TTCs containing short spacers show a state incompatible with NusG bridging and NusA binding (TTC-A; previously termed “expressome”). Structures of TTCs containing longer spacers reveal a new state compatible with NusG bridging and NusA binding (TTC-B) and reveal how NusG bridges and NusA binds. We propose that TTC-B mediates NusG- and NusA-dependent transcription-translation coupling.

**One Sentence Summary:** Cryo-EM defines states that mediate NusG- and NusA-dependent transcription-translation coupling in bacteria

## Main Text

Bacterial transcription and bacterial translation occur in the same cellular compartment, occur at the same time, and are coordinated processes, in which the rate of transcription by the RNA polymerase (RNAP) molecule synthesizing an mRNA is coordinated with the rate of translation by the first ribosome (“lead ribosome”) translating the mRNA (*1–9*; see, however, *10*). Data indicate that the coordination is mediated by transcription elongation factors of the NusG/RfaH family, which contain an N-terminal domain (N) that interacts with RNAP β’ and β subunits and a flexibly tethered C-terminal domain (C) that interacts with ribosomal protein S10, and which are thought to bridge, and thereby connect, the RNAP molecule and the lead ribosome (*2, 5–9*). Further data indicate that the coordination is modulated by the transcription elongation factor NusA (*11*).

Cramer and co-workers recently reported a 7.6 Å resolution cryo-EM structure of an *Escherichia coli* transcription-translation complex (TTC) termed the “expressome,” obtained by halting a transcription elongation complex (TEC) and allowing a translating ribosome to collide with the halted TEC (*12*). However, the mRNA molecule in the structure was not fully resolved, precluding determination of the number of mRNA nucleotides between the TEC and the ribosome active center in the structure (*12*), and the functional relevance of the structure has been challenged, due to its genesis as a collision complex, and due to its incompatibility with simultaneous interaction of NusG-N with RNAP and NusG-C with the ribosome (*6–9*). Demo *et al*. recently reported a ∼7 Å resolution cryo-EM structure of a complex of *E. coli* RNAP and a ribosome 30S subunit (*13*). However, the structure did not contain mRNA, did not position RNAP close to the 30S mRNA-entrance portal, and was incompatible with simultaneous interaction of NusG-N with RNAP and NusG-C with the ribosome (*13*).

Here, we report cryo-EM structures of *E. coli* TTCs containing defined-length mRNA spacers between the TEC and the ribosome active-center product site (P site), both in the presence of NusG and in the absence of NusG (Figs. 1, S1-S5; Tables S1-S2). We prepared synthetic nucleic-acid scaffolds that contained (i) DNA and mRNA determinants that direct formation of a TEC upon interaction with RNAP, (ii) an mRNA AUG codon that enables formation of a translation complex having the AUG codon positioned in the ribosome active-center P site upon interaction with a ribosome and tRNA^fMet^, and (iii) an mRNA spacer having a length, *n*, of 4, 5, 6, 7, 8, 9, or 10 codons (12, 15, 18, 21, 24, 27, or 30 nt) between (i) and (ii) (Fig. 1A). We then incubated the nucleic-acid scaffolds with RNAP, with ribosome and tRNA^fMet^, and optionally with NusG and/or NusA, and we determined structures by single-particle-reconstruction cryo-EM (see Supporting Information, Materials and Methods). With nucleic-acid scaffolds having short spacers (n = 4, 5, 6, 7, or 8), we obtained structures matching the “expressome” of *12* (TTC-A; Figs. 1B-left, 2, S1-S3; Table S1). However, with nucleic-acid scaffolds having longer mRNA spacers (n = 8, 9, or 10), we obtained structures of a new molecular assembly with features strongly suggesting it is the molecular assembly that functionally mediates NusG-dependent, NusA-dependent transcription-translation coupling in cells (TTC-B; Figs. 1B-center, 1B-right, 3-4, S3-S8; Table S1; Movies S1-S2).

**Fig. 1.**
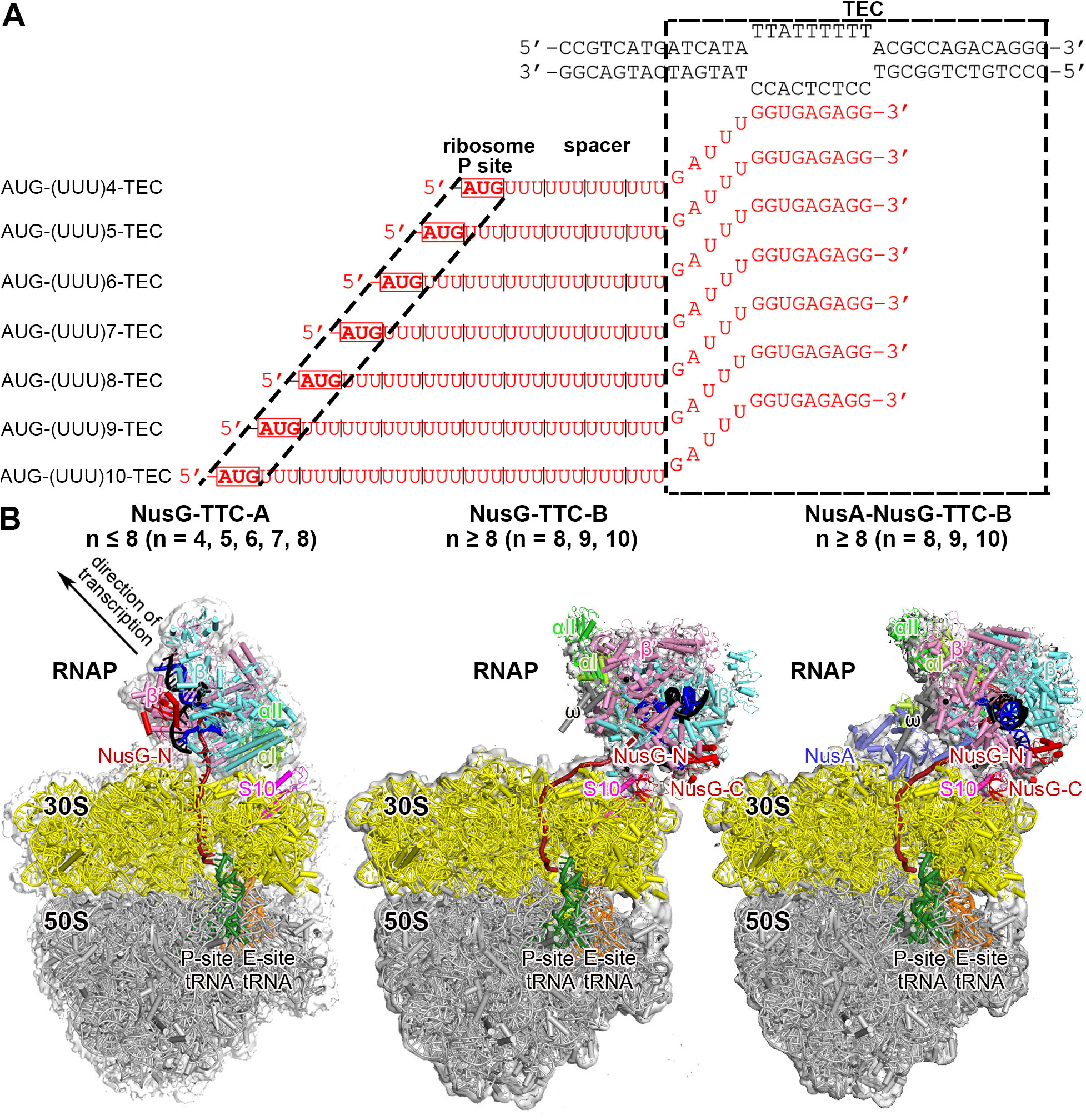
Structure determination: transcription-translation complexes (TTCs) **(A)** Nucleic-acid scaffolds. Each scaffold comprises nontemplate- and template-strand oligodeoxyribonucleotides (black) and one of seven oligoribonucleotides having spacer length n of 4, 5, 6, 7, 8, 9, or 10 codons (red), corresponding to mRNA. Dashed black box labeled “TEC,” portion of nucleic-acid scaffold that forms TEC upon addition of RNAP (10 nt nontemplate- and template-strand ssDNA segments forming “transcription bubble,” 10 nt of mRNA engaged with template-strand DNA as RNA-DNA “hybrid,” and 5 nt of mRNA, on diagonal, in RNAP RNA-exit channel); dashed black lines labeled “ribosome P-site,” mRNA AUG codon intended to occupy ribosome active-center P site upon addition of ribosome and tRNA^fMet^; “spacer,” mRNA spacer between TEC and AUG codon in ribosome active-center P site. **(B)** Cryo-EM structures of NusG-TTC-A (obtained with spacer lengths of 4-8 codons), NusG-TTC-B (obtained with spacer lengths of 8-10 codons), and NusA-NusG-TTC-B (obtained with spacer lengths of 8-10 codons). Structures shown are NusG-TTC-A (3.7 Å; n = 4; Table S1), NusG-TTC-B (4.7 Å; n = 9; Table S1), and NusA-NusG-TTC-B2 (3.5 Å; n = 8; Table S1). Images show EM density (gray surface) and fit (ribbons) for TEC, NusG and NusA (at top; direction of transcription, defined by downstream dsDNA, indicated by arrow in left panel and directly toward viewer in center and right panels) and for ribosome 30S and 50S subunits and P- and E-site tRNAs (at bottom). RNAP β’, β, α^I^, α^II^, and ω subunits are in pink, cyan, light green, and dark green, and gray; 30S subunit, 50S subunit, P-site tRNA, E-site tRNA are in yellow, gray, green, and orange; DNA nontemplate strand, DNA template strand, and mRNA are in black, blue, and brick-red (brick-red dashed line where modelled). NusG, NusA, and ribosomal protein S10 are in red, light blue, and magenta. Ribosome L7/L12 stalk omitted for clarity in this and all subsequent images.

**Fig. 2.**
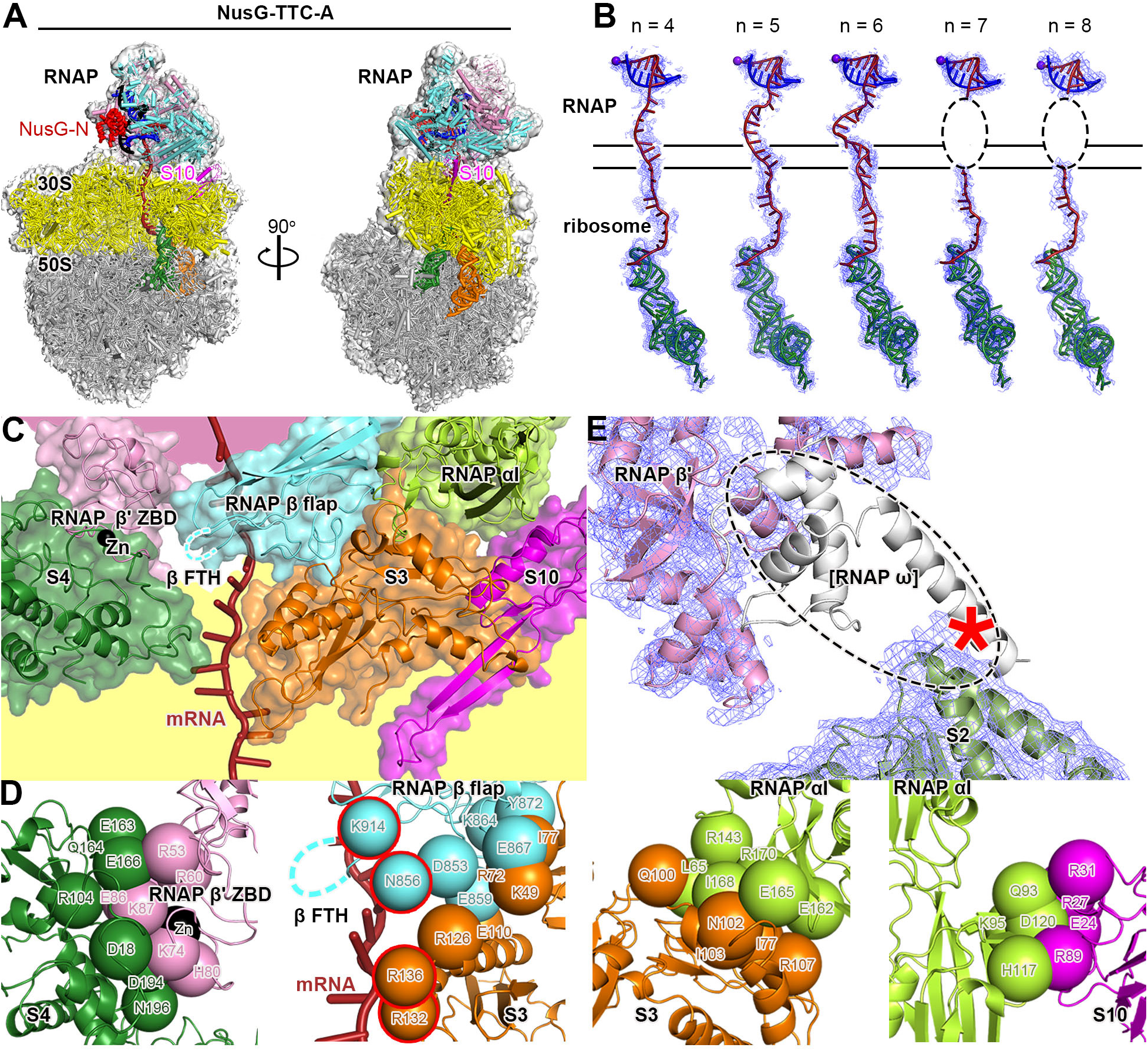
Cryo-EM structure of NusG-TTC-A. **(A)** Structure of NusG-TTC-A (3.7 Å; n = 4; Table S1). Two orthogonal views. Colors as in Fig. 1B. **(B)** Accommodation of mRNA spacer lengths of 4, 5, 6, 7 and 8 codons in NusG-TTC-A. EM density, blue mesh; mRNA, brick-red (disordered mRNA nucleotides indicated by dashed oval); template-strand DNA in RNA-DNA hybrid, blue; RNAP active-center catalytic Mg^2+^, purple sphere; tRNA in ribosome P site, green. Upper and lower black horizontal lines indicate edges of RNAP and ribosome. **(C)** RNAP-ribosome interface in NusG-TTC-A (n = 4; identical interface for n = 5, 6, 7, or 8), showing RNAP β’ zinc binding domain, (ZBD, pink; Zn^2+^ ion as black sphere), RNAP β flap, cyan, RNAP β flap tip helix (β FTH; disordered residues indicated by cyan dashed line), and RNAP α^I^ (green) interacting with ribosomal proteins S4 (forest green), S3 (orange), and S10 (magenta) and with mRNA (brick red). Portions of RNAP β’ and ribosome 30S not involved in interactions are shaded pink and yellow, respectively. **(D)** RNAP-ribosome interactions involving RNAP β’ ZBD and S4 (subpanel 1), RNAP β flap and S3 (subpanel 2; β FTH, dashed line; β and S3 residues that interact with mRNA, cyan and orange spheres with red outlines; mRNA, brick-red), RNAP α^I^ and S3 (subpanel 3), and RNAP α^I^ and S10 (subpanel 4). Other colors as in (C). **(E)** Absence of EM density for RNAP ω subunit. EM density, blue mesh; atomic models for RNAP β’ and S2, pink ribbon and forest-green ribbon, respectively; location of missing EM density for ω, dashed oval; ω in TEC in absence of ribosome (PDB 6P19; *17*), white ribbon.

**Fig. 3.**
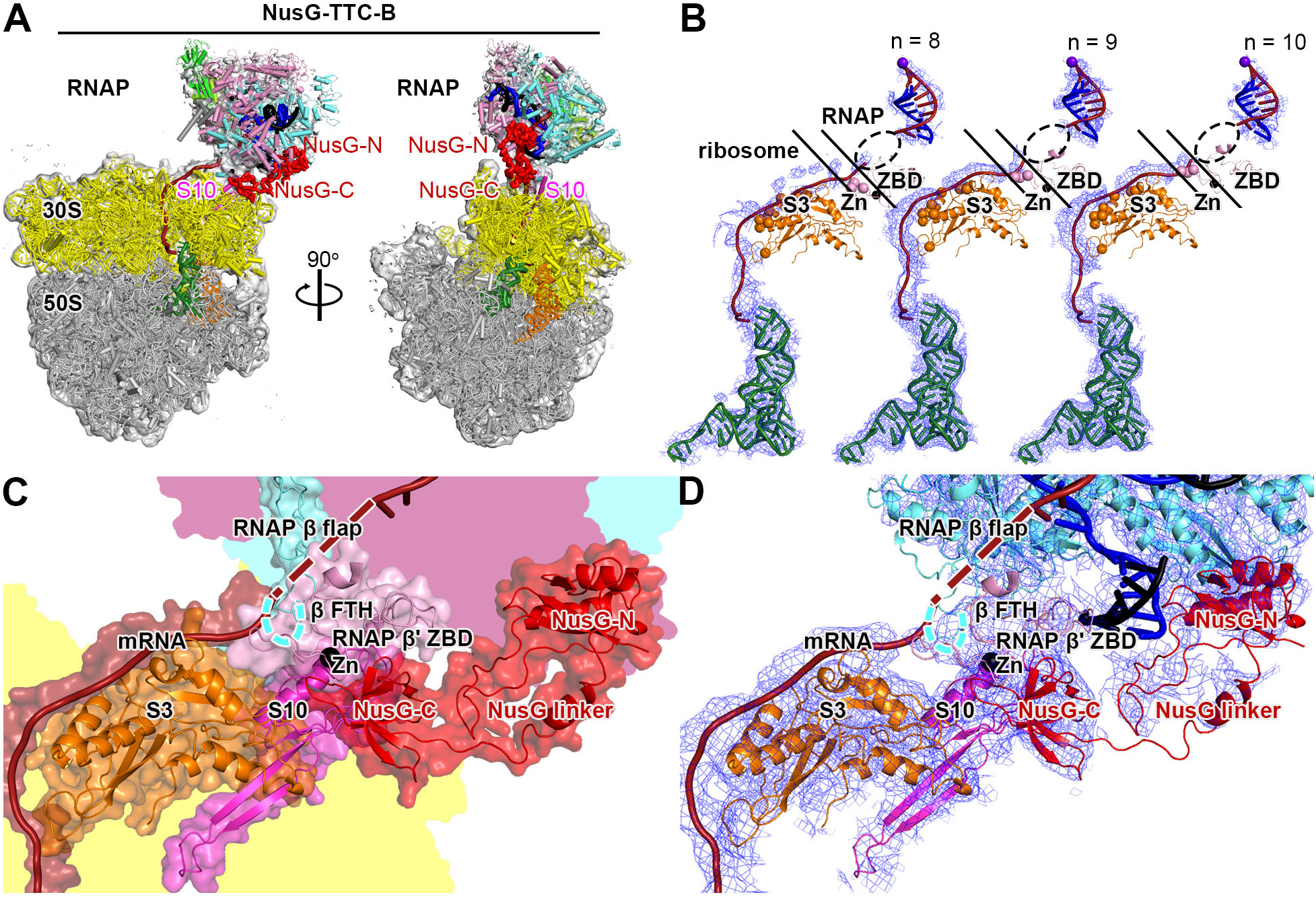
Cryo-EM structure of NusG-TTC-B. **(A)** Structure of NusG-TTC-B (4.7 Å; n = 9; Table S1). Views and colors as in Fig. 2A. **(B)** Accommodation of mRNA spacer lengths of 8, 9, and 10 codons in NusG-TTC-B. EM density, blue mesh; mRNA, brick-red (disordered mRNA nucleotides indicated by dashed oval); template-strand DNA in RNA-DNA hybrid, blue; RNAP active-center catalytic Mg^2+^, purple sphere; tRNA in ribosome P site, green; ribosomal protein S3, orange (positively charged residues positioned to contact mRNA as orange spheres); RNAP β’ zinc binding domain (ZBD, pink; Zn^2+^ ion as black sphere; positively charged residues positioned to contact mRNA as pink spheres). Upper and lower black diagonal lines indicate edges of RNAP and ribosome. **(C)** RNAP-ribosome interface and NusG bridging in NusG-TTC-B (n = 9; identical interface for n = 8, 9, or 10). RNAP β’ zinc binding domain, (ZBD, pink; Zn^2+^ ion as black sphere) interacts with ribosomal protein S3 (orange) and mRNA (brick red). NusG (red) bridges RNAP and ribosome, with NusG-N interacting with RNAP and NusG-C interacting with ribosomal protein S10 (magenta). Portions of RNAP β’, β, and ribosome 30S not involved in interactions are shaded pink, cyan, and yellow, respectively. **(D)** As C, showing cryo-EM density as blue mesh.

**Fig. 4.**
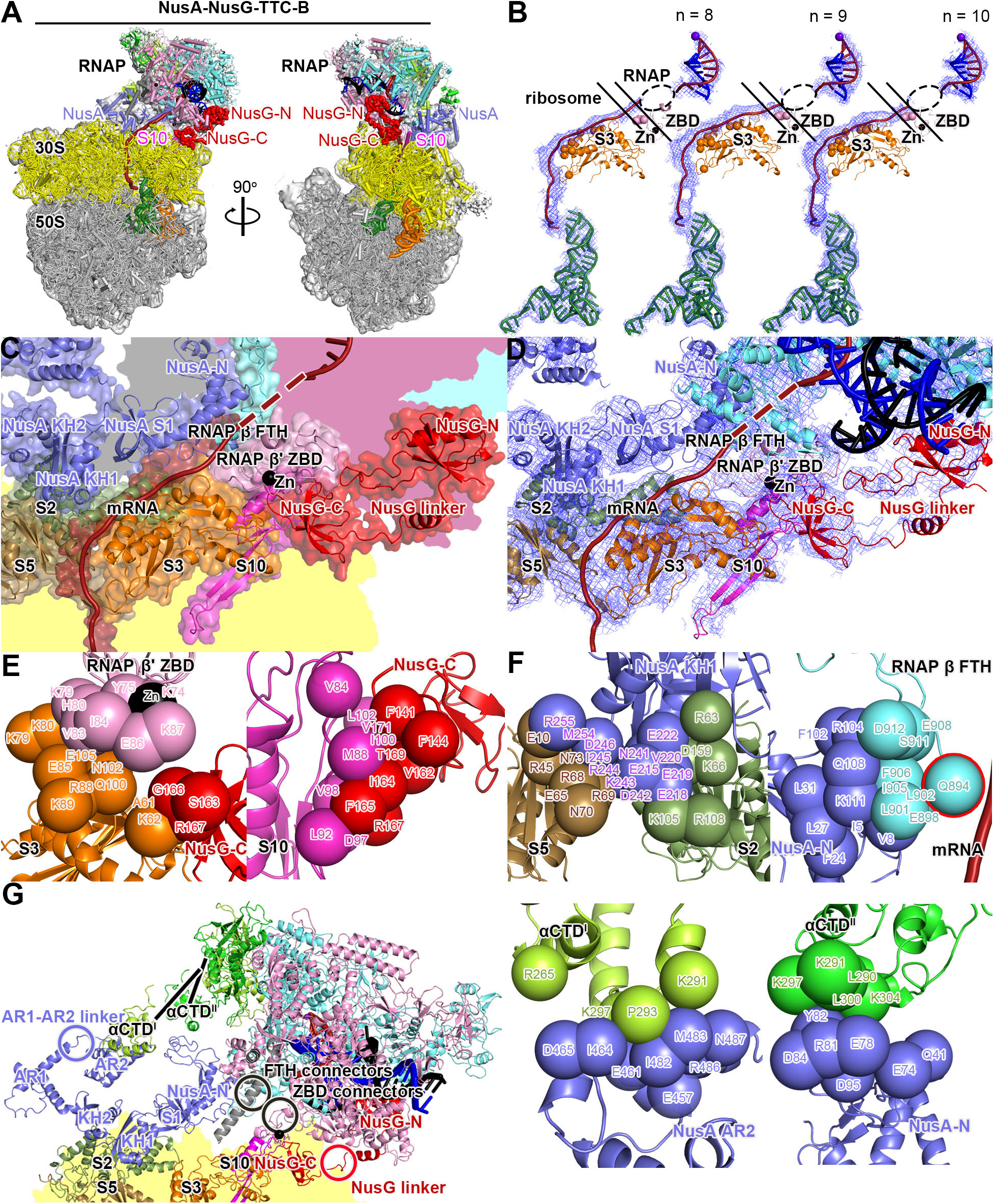
Cryo-EM structure of NusA-NusG-TTC-B. **(A)** Structure of NusA-NusG-TTC-B (NusA-NusG-TTC-B2; 3.5 Å; n = 9; Table S1). NusA, light blue. Views and other colors as in Figs. 2A and 3A. **(B)** Accommodation of mRNA spacer lengths of 8, 9, and 10 codons in NusA-NusG-TTC-B. Views and colors as in Fig 3B. **(C)** RNAP-ribosome interface, NusG bridging, and NusA binding in NusA-NusG-TTC-B (n = 9; identical interface for n = 8, 9, or 10). RNAP β’ zinc binding domain, (ZBD, pink; Zn^2+^ ion as black sphere) interacts with ribosomal protein S3 (orange) and mRNA (brick red). NusG (red) bridges RNAP and ribosome, with NusG-N interacting with RNAP and NusG-C interacting with ribosomal protein S10 (magenta). NusA (light blue) KH1 domain interacts with ribosomal proteins S5 and S2 (brown and forest green). Portions of RNAP β’, β, ω, and ribosome 30S not involved in interactions are shaded pink, cyan, gray, and yellow, respectively. **(D)** As C, showing cryo-EM density as blue mesh. **(E)** RNAP-ribosome interactions involving RNAP β’ ZBD and S3 (subpanel 1) and NusG-ribosome interactions involving NusG-C and S10 (subpanel 2). **(F)** NusA-ribosome interactions involving NusA KH1 and S5 and S2 (subpanel 1) and NusA-RNAP interactions involving NusA-N and RNAP β FTH (subpanel 2; β FTH residue that interacts with mRNA, cyan sphere with red outline; mRNA, brick-red), NusA AR2 and RNAP αCTD^I^ (subpanel 3), and NusA-N and RNAP αCTD^II^ (subpanel 4). **(G)** Points of flexibility in NusA-NusG-TTC-B (NusA “coupling pantograph”): flexible linkage in NusA structure (AR1-AR2 linker; light blue circle), three flexible linkages between NusA and RNAP (αCTD^I^ linker, αCTD^II^ linker, and β FTH connectors; black lines and black circle), flexible linkage between RNAP and ribosome (β’ ZBD connectors; black circle), and flexible NusG bridging of RNAP and ribosome (NusG linker; red circle).

TTC-A was obtained with nucleic-acid scaffolds having mRNA spacers of 4, 5, 6, 7, or 8 codons--but not with longer mRNA spacers (Figs. 1B-left, 2, S1-S3; Table S1). TTC-A was obtained both in the absence of NusG and in the presence of NusG (Figs. S1-S3; Table S1). EM density maps of 3.7-6.3 Å resolution were obtained (∼7 Å and ∼3.5 Å local resolution for TEC and ribosome, respectively, in best maps), enabling unambiguous rigid-body docking of atomic structures of TEC, ribosome 30S subunit, ribosome 50S subunit with tRNA in active-center P site and exit site (E site), and, where present, NusG-N, followed by manual fitting of residues in the RNAP-ribosome interface and in DNA and mRNA (Figs. 1B-left, 2, S1-S3; Table S1).

Remarkably, in TTC-A, the spatial relationship of RNAP relative to the ribosome is identical in structures obtained with nucleic-acid scaffolds having mRNA spacer lengths of 4, 5, 6, 7, and 8 codons (Fig. S1F). High-resolution data for TTC-A reveal that differences in mRNA spacer length are accommodated through differences in extents of compaction of mRNA in the RNAP RNA-exit channel and RNAP-ribosome interface (Fig. 2B). As mRNA spacer length increases from 4 codons to 5 codons to 6 codons, the number of mRNA nucleotides in the RNAP RNA-exit channel and RNAP-ribosome interface increases from 7 nt (5 nt in exit channel; 2 nt in interface) to 10 nt (7 nt in exit channel; 3 nt in interface) to 13 nt (9 nt in exit channel; 4 nt in interface) (Fig. 2B, subpanels 1-3). When the mRNA spacer length increases to 7 or 8 codons, 16 or 19 nt of mRNA are accommodated in the RNAP RNA-exit channel and RNAP-ribosome interface, and the 16 or 19 nt of mRNA show disorder, indicating they adopt an ensemble of different conformations (Fig. 2B, subpanels 4-5). We point out that the volume of the RNAP RNA-exit channel and RNAP-ribosome interface cannot accommodate more than ∼19 nt of mRNA without changing the conformation of the RNAP RNA-exit channel or disrupting the RNAP-ribosome interface, and we suggest that this accounts for our observations that TTC-A is obtained at relatively low particle populations with a nucleic-acid scaffold having an mRNA spacer length of 8 codons (18% vs. 91% for nucleic-acid scaffold having mRNA spacer length of 4 codons; Figs. S1, S3) and is not obtained with nucleic-acid scaffolds having mRNA spacer lengths >8 codons (Figs. S4-S5). The mRNA spacers analyzed in this work contained only U (Fig. 1A); because U is the RNA nucleotide having the smallest volume, the mRNA-spacer-length cut-off of 8 codons observed in this work is likely to represent an upper bound.

In TTC-A, the RNAP-ribosome interface is extensive (3,742 Å^2^ buried surface area) and involves contacts of RNAP β’ zinc-binding domain (ZBD), RNAP β flap, and RNAP α^I^ with ribosomal proteins S4, S3, and S10, respectively (Figs. 2C-D).

In EM density maps of TTC-A, density is absent for RNAP ω subunit, indicating that RNAP ω subunit is either absent, or at a low occupancy level, or disordered (Fig. 2E). Molecular modelling suggests that, if RNAP ω were present and fully folded, the C-terminal α-helix of ω would clash with the ribosome (Fig. 2E).

In EM density maps of TTC-A obtained in the presence of NusG, EM density is present for NusG-N (residues 1-118) at its expected binding location on the RNAP β’ clamp helices and the RNAP β pincer tip (*14*; Figs. 1B-left, S1H), but is absent for the NusG linker and NusG-C, consistent with unrestricted motion of the linker and NusG-C relative to NusG-N (*14*; Fig. 1B-left). Density maps for TTC-A obtained in the absence of NusG are identical to those obtained in the presence of NusG, except that density for NusG-N is missing (Fig. S2). Model building indicates that the shortest sterically allowed distance between NusG-N bound to RNAP and NusG-C modelled as bound to its molecular target on the ribosome, ribosomal protein S10 (*2, 5–9*), is 160 Å in TTC-A--which is 1.9 times the maximum length of the NusG linker--indicating that TTC-A is incompatible with NusG bridging of RNAP and S10 (Fig. S9A).

Molecular modelling indicates that TTC-A also is incompatible with other known functional properties of transcription elongation, pausing, and termination in *E. coli*. TTC-A is sterically incompatible with binding of NusA (*15*; Fig. S10A), formation of a 21 Q antitermination complex (*16–17*; Fig. S10B), and formation of pause and termination RNA hairpins (*15, 18–19*; Fig. S10C-D). TTC-A also appears to be incompatible with ribosome 30S-head swivelling, the 21° rotation of the ribosome 30S head relative to the ribosome 30S body that occurs during ribosome translocation (*20–22*; Fig. S11A; Movie S3). The RNAP-ribosome interface in TTC-A spans the 30S head and 30S body in the unswivelled state (Figs. 2C, S11A-left) and is expected to be disrupted upon swivelling (loss of 1,972 A^2^ buried surface area; Fig. S11A-right). The finding that TTC-A--the “expressome” of *12*--lacks RNAP ω subunit, is incompatible with NusG bridging, and is incompatible with known functional properties of transcription and translation in *E. coli* indicates that TTC-A is unlikely to be functionally relevant to transcription-translation coupling under most conditions in *E. coli*. We propose that TTC-A is either: (i) a specialized complex that mediates transcription-translation coupling under specialized circumstances (e.g., transcription-translation coupling by RNAP deficient in ω or by ribosomes inactive in translocation), or an anomalous complex formed when the mRNA spacer between RNAP and ribosome is anomalously short (e.g. “collision-ome” or “crash-ome”).

TTC-B was obtained with nucleic-acid scaffolds having mRNA spacer lengths of 8, 9, or 10 codons--but not with shorter mRNA spacers (Figs. 1B, 3-4, S3-S7; Table S1). TTC-B was obtained only when NusG was present (Figs. S3-S8; Table S1) and was obtained both without bound NusA and with bound NusA (Figs. S3-S7; Table S1). TTC-B differs from TTC-A by translation of RNAP relative to the ribosome by ∼70 Å and rotation of RNAP relative to the ribosome by ∼180° (Fig. 1B; Movie S1). EM density maps at 3.1-12.6 Å resolution were obtained, (∼7 Å and ∼3 Å local resolution for TEC and ribosome, respectively, in best maps), enabling unambiguous rigid-body docking of atomic structures of components, followed by manual fitting (Figs. 3-4, S3-S7). TTC-B is identical to the NusG-bridged complex reported in a preprint by Weixlbaumer and co-workers (*23*) and is different from the NusA-containing complex reported in a preprint by Mahamid, Rappsilber, and co-workers (*24*).

In contrast to in TTC-A, where the RNAP RNA-exit channel is coupled directly to the ribosome mRNA entrance portal, in TTC-B, the RNAP RNA-exit channel is separated by ∼60 Å from the ribosome mRNA entrance portal (Fig. 1B). In TTC-B, a ∼60 Å, ∼11 nt, mRNA segment connects the RNAP RNA-exit channel and the ribosome mRNA entry portal, running along the surface of the ribosome 30S head, making favorable electrostatic interactions with positively charged residues in ribosomal protein S3 and RNAP β’ ZBD (Figs. 3B, 4B, S3F, S4G). The requirement for this additional ∼11 nt mRNA segment accounts for the fact that TTC-B is obtained only with nucleic-acid scaffolds having mRNA spacer lengths ≥8 codons.

In TTC-B, the spatial relationship of RNAP relative to the ribosome is identical in structures obtained with mRNA spacer lengths of 8, 9, and 10 codons (Figs. S4F, S5G). Analogously to in TTC-A, in TTC-B, differences in mRNA spacer length are accommodated through differences in extents of compaction of mRNA in the RNAP RNA-exit channel (Figs. 3B, 4B). As mRNA spacer length increases from 8 to 9 to 10 codons, the number of mRNA nucleotides in the RNAP RNA-exit channel increases from ∼8 nt to ∼11 nt to ∼14 nt (disordered in each case; Figs. 3B, 4B). Assuming that the volume of the RNAP RNA-exit channel allows it to accommodate up to ∼15 nt of mRNA (see above), it seems likely that mRNA spacer lengths up to ∼10 codons could be accommodated in TTC-B. Noting that the mRNA segments in the RNAP-ribosome interface and near ribosomal protein S3 in TTC-B are solvent-accessible, it also seems possible that longer, possibly much longer, mRNA spacer lengths could be accommodated by looping of, or secondary-structure formation in, these mRNA segments.

In TTC-B, the interaction between RNAP and ribosome is small, involving only contact between the RNAP β’ ZBD sequence and ribosomal protein S3 (224 Å^2^ buried surface area; Figs. 3C-D, 4C-E). In TTC-B, the RNAP-ribosome interaction is supplemented by bridging of RNAP and the ribosome by NusG, involving simultaneous binding of NusG-N to RNAP and binding of NusG-C to ribosomal protein S10 (1,409 Å^2^ buried surface area for NusG-C and S10; Figs. 1B, 3A,C-D, 4A,C-E, S3G, S4H, S9B-C). NusG-C interacts with S10 in the manner expected from published structures of a complex of NusG and S10 and of a complex of NusG and a ribosome (*2*, 9; Figs. S3G, S4H). EM density maps show unambiguous density for NusG-N, NusG-C, and most residues of the NusG linker (Figs. 3D, 4D, S3G, S4H), and, at lower contour levels, show density for all residues of the NusG linker (Figs. 3C, 4C). Corresponding EM maps obtained in the absence of NusG do not show TTC-B (Fig. S8), indicating that NusG bridging is functionally important for the formation and/or the stability of TTC-B. The NusG bridging hypothesized in refs. *2* and *9* is unequivocally verified.

We first obtained structures of TTC-B in the presence of NusG and absence of NusA (NusG-TTC-B; Figs. 3, S3-S4; Table S1). Molecular modelling indicated that NusG-TTC-B potentially could accommodate binding of NusA (Fig. S10A). Therefore, we sought, and obtained, corresponding structures of TTC-B in the presence of *both* NusG and NusA (NusA-NusG-TTC-B; Figs. 4, S5; Table S1). As compared to structure determination of TTC-B in the absence of NusA, structure determination of TTC-B in the presence of NusA was associated with substantially higher particle populations (4% vs. 45% for n = 8, 18% vs. 28% for n = 9, and 17% vs. 40% for n = 10) and substantially higher resolutions (12.6 Å vs. 3.1 Å for n = 8, 4.7 Å vs. 4.2 Å for n = 9, and 5.0 Å vs. 3.7 Å for n = 10), indicating that NusA functionally stabilizes TTC-B. Three NusA-NusG-TTC-B subclasses were obtained: TTC-B1, TTC-B2, and TTC-B3, differing by up to 15° rotation of RNAP relative to NusA and ribosome (Figs. S5, S7A-B; Movie S2).

In all NusA-NusG-TTC-B subclasses, RNAP and NusG interact with the ribosome 30S head, with RNAP β’ ZBD contacting ribosomal protein S3 and NusG contacting ribosomal protein S10 (Fig. 4C-E, S6), essentially as in the absence of NusA (Fig. 3C-D).

In all NusA-NusG-TTC-B subclasses, NusA makes identical--and extensive--interactions with the surface of the ribosome S30 body, involving contacts between NusA KH1 domain and ribosomal proteins S2 and S5 (1,755 Å^2^ buried surface area; Figs. 4C-E, S6). The NusA-ribosome interactions observed here show no similarity to the putative NusA-ribosome interactions reported in *24*; the orientation of NusA relative to the ribosome differs by ∼180°, and the interactions involve a different module of the ribosome 30S subunit (body vs. head).

NusA functions in this context as a large--70 Å x 50 Å--open rectangular frame that connects RNAP to the ribosome 30S body (Figs. 4G, S7C). One side of the NusA rectangular frame interacts with the ribosome 30S body, and three corners of the NusA rectangular frame interact with RNAP, contacting the RNA α^I^ C-terminal domain (αCTD^I^), the RNA α^II^ C-terminal domain (αCTD^II^), and the RNAP β flap-tip helix (FTH) (Figs 4F-G, S7C). The NusA rectangular frame contains an internal flexible linkage, the AR1-AR2 linker (light blue circle in Figs. 4G, S7C), and interacts with RNAP through three flexible linked modules: αCTD^I^ and αCTD^II^, which are connected to the rest of RNAP through long, flexible linkers (*25*; lines in Figs. 4G, S7C), and β FTH, which is connected to the rest of RNAP through flexible connectors (*15–18*; black circle in Figs. 4G, S7C). The internal flexibility and flexible connections enable the NusA-RNAP subcomplex to maintain constant contact with the ribosome 30S body, despite differences in orientation of RNAP relative to the ribosome 30S body (Fig. S7C; Movie S2). We refer to the NusA rectangular frame as the “coupling pantograph,” analogizing it to an electric-railway coupling pantograph, the open rectangular frame, with internal flexibility and flexible connections, that enables a locomotive to maintain constant contact with a power cable, despite differences in orientation of the locomotive relative to the cable (*26*; Figs. S7C; Movie S2).

The separation between the RNAP RNA-exit channel and the ribosome mRNA entry portal in TTC-B, together with the open character of the NusA rectangular frame (“coupling pantograph”) in TTC-B, provides largely unrestricted access for transcriptional-regulatory factors to bind, and transcriptional-regulatory RNA secondary structures to form, at and adjacent to the mouth of the RNAP RNA-exit channel (Fig. S10). Molecular modelling indicates that TTC-B, unlike TTC-A, can accommodate formation of the 21 Q antitermination complex (*16–17*; Fig. S10B) and can accommodate formation of pause and termination RNA hairpins (*15, 18–19*; Fig. S10C-D). In NusA-NusG-TTC-B, positively charged residues of NusA N and S1 domains are positioned to make favorable electrostatic interactions with the hairpin loop of a pause or termination RNA hairpin, and thereby potentially to nucleate formation of a pause or termination RNA hairpin (Fig. S7D; see *15*). The different orientations of NusA N and S1 domains in NusA-NusG-TTC-B subclasses B1, B2, and B3 possibly enable interactions with different-length pause and termination RNA hairpins, with B1 accommodating shorter hairpins and B2 and B3 accommodating longer hairpins (Fig. S7D).

Molecular modelling also indicates that TTC-B, unlike TTC-A, is compatible with ribosome 30S-head swivelling, the rotation of the 30S head relative to the 30S body that occurs during ribosome translocation (*20–22*; Fig. S11B-C; Movies S4-S5). In NusG-TTC-B, all RNAP-ribosome and NusG-ribosome interactions involve the ribosome 30S head; accordingly, 30S-head swivelling can be accommodated by rotation of RNAP and NusG with the 30S head (Fig S11B, center) and/or by separate rotation of flexibly connected RNAP β’ ZBD and flexibly connected NusG-C with the 30S head (Fig S11B, right; Movie S4). In NusA-NusG-TTC-B, NusA-ribosome interactions involve the ribosome 30S body, and RNAP-ribosome and NusG-ribosome interactions involve the ribosome 30S head; nevertheless--exploiting the internal flexibility and flexible connections of the NusA-RNAP “coupling pantograph”--30S-head swivelling can be accommodated by rotation of RNAP and NusG with the 30S head (Fig S11B, center) and/or by separate rotation of flexibly connected RNAP β’ ZBD and flexibly connected NusG-C with the 30S head (Fig S11B, right; Movie S5).

Based on the observation that TTC-B is compatible with NusG bridging, NusA binding, known functional aspects of transcription, and known functional aspects of translation, we propose that TTC-B modulates NusG-dependent, NusA-dependent transcription-translation coupling in *E. coli*.

The structures presented were determined in the presence of CHAPSO, a non-ionic detergent that has been used extensively in cryo-EM structural analysis of RNAP and RNAP complexes to improve structural homogeneity by disrupting non-specific complexes and weak complexes, and by improving rotational-orientation distributions of particles by reducing interactions with the air-water interface (see *14–18*). Analogous structure determination in the absence of CHAPSO yielded low-resolution maps of TTC-A for nucleic-acid scaffolds with mRNA spacer lengths of 4, 5, 6, and 7 codons (Figs. S12-S13; Tables S2; Movie S6) and yielded low-resolution maps of two additional complexes, TTC-C and TTC-D, for nucleic-acid scaffolds with mRNA spacer lengths of 7, 8, and 9 codons (Figs. S13-S16; Table S2; Movies S7-S11). The fact that TTC-C and TTC-D are observed only in the absence of CHAPSO suggests TTC-C and TTC-D may involve relatively weak interactions. In TTC-C and TTC-D, interactions between RNAP and ribosome are mediated by RNAP β sequence insert 2 (βSI2; also known as βi9; *26–27*), a 60-Å long α-helical antiparallel coiled-coil flexibly tethered to the rest of RNAP, and the main interaction is an electrostatic interaction between the tip of βSI2 and the ribosome 30 S subunit (Figs. S15-S16). In TTC-C the orientation of RNAP relative to the ribosome is compatible with NusG bridging (Fig S15), and in TTC-D the orientation of RNAP relative to the ribosome is incompatible with NusG bridging (Fig S16). The structures suggest that TTC-C and TTC-D could play roles in NusG-dependent transcription-translation coupling and in NusG-independent transcription-translation coupling, respectively. The structural module that mediates RNAP-ribosome interaction in TTC-C and TTC-D--βSI2--is not essential for growth in rich media (*26*), but is essential for growth in minimal media (*26*), implying that TTC-C and TTC-D are unlikely to be important for transcription-translation coupling in general, but may be important for transcription-translation coupling in specific transcription units in specific regulatory contexts (see *6*). Further analysis will be needed to determine whether, and, if so, in which contexts, TTC-C and TTC-D function in transcription-translation coupling in *E. coli.* The results presented define four structural classes of TTCs--TTC-A (the previously reported “expressome”; *12*), TTC-B, TTC-C, and TTC-D--and show that TTC-B has structural properties indicating it mediates NusG-dependent, NusA-dependent transcription-translation coupling in *E. coli*.

The results presented reframe our understanding of the structural and mechanistic basis of transcription-translation coupling. The results provide high-resolution structures of the previously described “expressome” (*12*; TTC-A) that demonstrate the incompatibility of the previously described “expressome” with general transcription-translation coupling. In addition, the results provide high-resolution structures of a new structural state, TTC-B, with properties assignable to general, NusG-dependent, NusA-dependent transcription-translation coupling, show that NusG stabilizes TTC-B by bridging RNAP and the ribosome 30S head, show that NusA stabilizes TTC-B by bridging RNAP and the ribosome 30S body, and show that NusA serves as a “coupling pantograph” that bridges RNAP and the ribosome 30S body in a flexible manner that allows rotation of RNAP relative to the ribosome 30S body. Finally, the results provide testable new hypotheses regarding the identities of the RNAP and NusA structural modules crucial for transcription-translation coupling (RNAP β’ ZBD and NusA KH1) and regarding the interactions made by those structural modules (interactions with ribosomal protein S3 in the S30 head and interactions with ribosomal proteins S2 and S5 in the S30 body).

## Acknowledgements

We thank the Rutgers University Cryo-EM Core facility, University of Michigan Life Sciences Institute Cryo-EM Facility, National Center for CryoEM Access and Training (supported by NIH grant GM129539, Simons Foundation grant SF349247, and New York state grants), and Pacific Northwest Center for Cryo-EM (supported by NIH grant GM129547 and Department of Energy Environmental Molecular Sciences Laboratory) for microscope access; K. Kuznedelov and K. Severinov for plasmids; and L. Minakhin, B. Nickels, and J. Winkelman for discussion; and E. Eng and H. Wei for assistance.

## Funding

This work was supported by University of California discretionary funds to G.B., University of Michigan discretionary funds to M.S., and National Institutes of Health (NIH) grant GM041376 to R.H.E.

## Author Contributions

V.M. and G.B. prepared biomolecules. C.W., V.M., E.F., J.K., and M.S. collected data. C.W., V.M, J.K., M.S., and R.H.E. analyzed data. C.W., V.M., and R.H.E. prepared figures. R.H.E. designed experiments and wrote the manuscript.

## Data and Material Availability

Cryo-EM micrographs have been deposited in the Electron Microscopy Public Image Archive Resource (EMPIAR accession codes 10467 and 10468). Cryo-EM maps and atomic models have been deposited in the Electron Microscopy Database (EMDB accession codes 21386, 21468, 21469, 21470, 21471, 21472, 21474, 21475, 21476, 21477, 21482, 21483, 21485, 21486, 21494, 22082, 22084, 22087, 22107, 22141, 22142, 22181, 22192, and 22193) and the Protein Database (PDB accession codes 6VU3, 6VYQ, 6VYR, 6VYS, 6VYT, 6VYU, 6VYW, 6VYX, 6VYY, 6VYZ, 6VZJ, 6VZ2, 6VZ3, 6VZ5, 6VZ7, 6XDQ, 6XDR, 6XGF, 6XII, 6XIJ, 6X6T, 6X7F, 6X7K, and 6X9Q).

## Supplementary Materials

### Materials and Methods

#### *E. coli* RNAP core enzyme

*E. coli* RNAP core enzyme was prepared from *E. coli* strain BL21 Star (DE3) (ThermoFisher) transformed with plasmid pVS10 (*29*; encodes *E. coli* RNAP β’ with C-terminal hexahistidine tag, β, α, and ω subunits), as described (*30*). The product (purity >95%) was stored in aliquots in RNAP storage buffer (10 mM Tris-HCl, pH 7.6, 100 mM NaCl, 0.1 mM EDTA, and 5 mM dithiothreitol) at −80°C.

#### *E. coli* NusA

*E. coli* NusA was prepared from *E. coli* strain BL21 Star (DE3) (ThermoFisher) transformed with plasmid pET28a-NusA (encodes *E. coli* NusA with N-terminal hexahistidine tag; *31*), as described (*31*). The product (purity >95%) was stored in aliquots in RNAP storage buffer at −80°C.

#### *E. coli* NusG

*E. coli* NusG was prepared from *E. coli* strain BL21 Star (DE3) (ThermoFisher) transformed with plasmid pIA247 (*32*; K. Kuznedelov and K. Severinov; encodes *E. coli* NusG with C-terminal hexahistidine tag), as described (*33*). The product (purity >95%) was stored in aliquots in RNAP storage buffer at −80°C.

#### *E. coli* 70S ribosome

*E. coli* 70S ribosomes were prepared from *E. coli* strain JE28 (*34*; grown in media containing 50 μg/ml kanamycin to OD600 = 1.0 in a 50 L fermenter, harvested, flash frozen in liquid N_2_, and stored at −80°C), as described (*35–36*).

Cells in 20 mM Tris-HCl, pH 7.6, 100 mM NH_4_Cl, 10.5 mM Mg(OAc)_2_, 0.5 mM EDTA, 4 mM 2-mercaptoethanol, and 1.5 mg/ml benzamidine HCl, and phenylmethylsulfonyl fluoride at 4°C were lysed [EmulsiFlex C3 (Avestin); 3 passages at 8,000 psi], and lysates were cleared by centrifugation (SS-34 rotor; 2 x 30 min at 16,000 rpm at 4°C). Ribosomes were pelleted by centrifugation through a 12 ml high-salt sucrose cushion [20 mM Tris-HCl, pH 7.5, 1.1 M sucrose, 500 mM NH_4_Cl, 10.5 mM Mg(OAc)_2_, 0.5 mM EDTA, and 4 mM 2-mercaptoethanol; Type 45 Ti rotor; 20 h at 43,000 rpm at 4°C; *37*]. Pelleted ribosomes were dissociated into ribosome 30S and 50S subunits by re-suspension in dissociation buffer [20 mM Tris-HCl, pH 7.6, 200 mM NH_4_Cl, 1 mM Mg(OAc)_2_, and 4 mM 2-mercaptoethanol], and 30S and 50S subunits were separated by loading an aliquot (250 A_260_ units) onto a 36 ml 10-30% sucrose gradient in dissociation buffer and centrifugation (SW28 rotor; 15 h at 20,000 rpm at 4°C). The gradient was fractionized, fractions containing 30S and 50S subunits were identified by UV absorbance and pooled, and pooled fractions containing 30S and 50S subunits were pelleted (Type 45 rotor; 21 h at 41,000 rpm at 4°C). Purified 30S and 50S subunits were reassociated by suspension in reassociation buffer (20 mM Tris-HCl, pH 7.5, 30 mM KCl, 20 mM Mg(OAc)_2_, and 4 mM 2-mercaptoethanol), dilution to 200 A_260_ units/ml, and incubation 30 min at 40°C, and 70S ribosomes were separated from non-reassociated 30S and 50S subunits by loading a 1 ml aliquot onto a 36 ml 10-30% sucrose gradient in reassociation buffer and centrifugation (SW28 rotor; 15 h at 18,000 rpm at 4°C). The gradient was fractionized, fractions containing 70S ribosomes were identified by UV absorbance and pooled, and pooled fractions were pelleted (Type 45 rotor; 21 h at 41,000 rpm at 4°C). Pelleted 70S ribosomes were re-suspended in reassociation buffer, flash frozen in liquid N_2_, and stored in aliquots at −80°C.

#### *E. coli* tRNA^fMet^

*E. coli* tRNA^fMet^ was purchased (MP Biomedical), dissolved to 100 μM in 5 mM Tris-HCl, pH 7.5, and stored in aliquots at −80°C.

#### Nucleic-acid scaffolds

Oligodeoxyribonucleotides and oligoribonucleotides (sequences in Fig. 1A) were purchased (Integrated DNA Technologies; PAGE-purified), dissolved to 1 mM in 5 mM Tris-HCl, pH 7.5, and stored in aliquots at-80°C. Nucleic-acid scaffolds (sequences in Fig. 1A) were prepared by mixing 60 μM nontemplate-strand oligodeoxyribonucleotide, 60 μM template-strand oligodeoxyribonucleotide, and 60 μM oligoribonucleotide in 100 µl annealing buffer (5 mM Tris-HCl, pH 7.5), heating 10 min at 95°C, and cooling slowly (3 h) to 22°, and nucleic-acid scaffolds were stored in aliquots at −80°C.

#### Transcription-translation complexes (TTCs)

TTCs for structure determination in the presence of CHAPSO were prepared by mixing 10 µl 20 µM RNAP core enzyme (in RNAP storage buffer), 0 or 30 μl 30 μM NusG (in RNAP storage buffer), 0 or 16 μl 60 μM NusA (in RNAP storage buffer), 4 µl 60 µM nucleic-acid scaffold (in 5 mM Tris-HCl, pH 7.5), 10 μl 10x reassociation buffer, and 10 μl (reactions with both NusA and NusG), 26 μl (reactions with NusG but not NusA), or 40 μl (reactions with NusA but not NusG), or 56 μl (reactions with neither NusA nor NusG) water; incubating 10 min at 22°C; adding 10 µl 100 µM tRNA^fMet^ (in 5 mM Tris-HCl, pH 7.5), 5 µl 40 µM 70S ribosome (in reassociation buffer), and 5 μl water; and further incubating 10 min at 22°C. The resulting TTCs were transferred to a pre-chilled 0.5 ml Amicon Ultracel 30K concentrator (EMD Millipore), concentrated to 35 µl by centrifugation (15 min at 20,000xg at 4°C), incubated 30 min on ice for 30 min, supplemented with 3.8 µl of ice-cold 80 mM CHAPSO, and immediately applied to grids (see “Cryo-EM structure determination: sample preparation”).

TTCs for structure determination in the absence of CHAPSO were prepared by mixing 1 µl 20 µM RNAP core enzyme (in RNAP storage buffer), 3 µl 30 µM NusG (in RNAP storage buffer) and 0.4 µl 60 µM nucleic-acid scaffold (in 5 mM Tris-HCl, pH 7.5), 1 μl 10x reassociation buffer, and 2.6 μl water; incubating 10 min at 22°C; adding 1 µl of 100 µM tRNA^fMet^ (in 5 mM Tris-HCl, pH 7.5), 0.5 µl 40 µM 70S ribosomes (in reassociation buffer), and 0.5 μl water; and further incubated 10 min at 22°C. The resulting TTCs were incubated 30 min on ice, mixed with 35 µl ice-cold reassociation buffer, and immediately applied to grids (see “Cryo-EM structure determination: sample preparation”).

#### Cryo-EM structure determination: sample preparation

EM grids were prepared using a Vitrobot Mark IV autoplunger (FEI/ThermoFisher), with the environmental chamber at 22°C and 100% relative humidity. Samples (3 μl) were applied to 2/1 Quantifoil Cu 300 holey-carbon grids (Quantifoil); glow-discharged 60 s using a PELCO glow-discharge system (Ted Pella)]), grids were blotted with #595 filter paper (Ted Pella) for 7 s at 22°C, and grids were flash-frozen by plunging in a liquid ethane cooled with liquid N_2_, and grids were stored in liquid N_2_.

#### Cryo-EM structure determination: data collection and data reduction (NusG-TTC-A; n = 4, 5, 6, or 7; with CHAPSO)

Cryo-EM data for NusG-TTC-A (n = 5; with CHAPSO) were collected at the Rutgers University Cryo-EM core Facility, using a 200 kV Talos Arctica (FEI/ThermoFisher) electron microscope equipped with a GIF Quantum K2 direct electron detector (Gatan). Data were collected automatically in counting mode, using EPU (FEI/ThermoFisher), a nominal magnification of 130,000x, a calibrated pixel size of 1.038 Å/pixel, and a dose rate of 4.8 electrons/pixel/s. Movies were recorded at 200 ms/frame for 6 s (30 frames), resulting in a total radiation dose of 26.7 electrons/Å^2^. Defocus range was varied between −1.25 µm and −2 µm. A total of 1,027 micrographs were recorded from one grid over two days. Micrographs were gain-normalized and defect-corrected.

Data were processed as summarized in Figs. S1A-D. Data processing was performed using a Tensor TS4 Linux GPU workstation with four GTX 1080 Ti graphic cards (NVIDIA). Dose weighting motion correction (3×3 tiles; b-factor = 150) were performed using Motioncor2 (*37*). Contrast-transfer-function (CTF) estimation was performed using CTFFIND-4.1 (*38*). Subsequent image processing was performed using Relion 3.0 (*39*). Automatic particle picking with Laplacian-of-Gaussian filtering yielded an initial set of 98,720 particles. Particles were extracted into 500×500 pixel boxes and subjected to rounds of reference-free 2D classification and removal of poorly populated classes, yielding a selected set of 27,378 particles. The selected set was 3D-classified with C1 symmetry, using a *de novo* 3D template created using 3D_initial_model under Relion 3.0. Of five classes obtained, four classes exhibited strong, well-defined density assignable to the TEC and having a spatial relationship to density for the ribosome consistent with a TTC. The 24,959 particles for these four classes were combined and 3D auto-refined using a mask with a diameter of 450 Å. The resulting 3D auto-refined particles were further refined using a soft mask and solvent flattening and were post-processed, yielding a reconstruction at 3.7 Å overall resolution, as determined from gold-standard Fourier shell correlation (FSC; Fig. S1D; Table S1).

The initial atomic model for NusG-TTC-A (n = 5; with CHAPSO) was built by manual docking of RNAP β’, β, a^I^, a^II^ segments and NusG segments from a cryo-EM structure of an *E. coli* TEC bound to NusG-N (PDB 6C6U; *40*); DNA and RNA segments from a cryo-EM structure of an *E. coli* 21Q transcription antitermination “loaded” complex (PDB 6P19; *17*); ribosome 30S S2-S21 segments and 16S rRNA segments from a cryo-EM structure of an *E. coli* “expressome” (PDB 5MY1; *12*); ribosome 30S S1 segments from cryo-EM structure of an *E. coli* ribosome complex (PDB 6H4N, *41*); ribosome 50S subunit from a cryo-EM structure of an *E. coli* 50S ribosomal subunit (PDB 6QDW); ribosome 50S-stalk L7/12, L10, and L11 segments from a cryo-EM structure of an *E. coli* ribosome complex (PDB 6I0Y, *42*), and P- and E-site tRNA segments from a crystal structure of a *Thermus thermophilus* 70S ribosome P- and E-site tRNA and mRNA (PDB 4V6G, *43*), using UCSF Chimera (*44*). For the RNAP β’ N and C-termini (residues 1-15 and 1734-1407), the RNAP β flap-tip helix (residues 892-910), the RNAP α^I^ and α^II^ N-termini and C-terminal domain (residues 1-5 and 234-329), and the NusG linker and NusG-C (residues 117-182), density was absent, suggesting high segmental flexibility; these segments were not fitted. For the RNAP ω, density was absent, suggesting absence, low occupancy, or high segmental flexibility; RNAP ω was not fitted.

Refinement of the initial model was performed using real_space_refine under Phenix (*45*). The ribosome 30S and 50S subunits were rigid-body refined against the map, followed by real-space refinement with geometry, rotamer, Ramachandran-plot, Cβ, non-crystallographic-symmetry, secondary-structure, and reference-structure (initial model as reference) restraints, followed by global minimization and local-rotamer fitting. Secondary-structure annotation was inspected and edited using UCSF Chimera. RNAP and NusG were rigid-body refined against the map, and the RNAP β’ Zn^2+^ binding domain (ZBD; residues 41-100), RNAP β flap (residues 835-891 and 911-937), RNAP α^I^ (residues 61-75 and 154-173), DNA, and mRNA segments were subjected to iterative cycles of model building and refinement in Coot (*46*). The final atomic model at was deposited in the Electron Microscopy Data Bank (EMDB) and the Protein Data Bank (PDB) with accession codes EMDB 21468 and PDB 6VYQ (Table S1).

The cryo-EM structure of NusG-TTC-A (n = 4; with CHAPSO) was determined in the same manner, but using data collected at the National Center for CryoEM Access and Training (NCCAT), using a 300 kV Krios Titan (FEI/ThermoFisher) electron microscope equipped with a Gatan K2 Summit direct electron detector (Gatan), Leginon (*47*), a nominal magnification of 105,000x, a calibrated pixel size of 1.096 Å/pixel, a dose rate of 6.33 electrons/pixel/s and 200 ms/frame for 10 s (50 frames; total radiation dose of 63.5 electrons/Å^2^), and a defocus range of 1 µm to 3 µm, and recording a total of 6,724 micrographs were recorded from one grid over 1.5 days. The final map and atomic model had resolution of 3.8 Å (EMDB 21386; PDB 6VU3; Table S1; Fig. S1F).

Cryo-EM structures of NusG-TTC-A (n = 6; with CHAPSO) and NusG-TTC-A (n = 7; with CHAPSO) were determined in the same manner as for NusG-TTC-A (n = 5; with CHAPSO), yielding maps and atomic models with resolutions of 3.8 Å (EMDB 21469; PDB 6VYR) and 3.7 Å (EMDB 21470; PDB 6VYS), respectively (Table S1; Fig S1F).

The cryo-EM structure of TTC-A (n = 5; with CHAPSO) in the absence of NusG was determined in the same manner as for NusG-TTC-A (n = 5; with CHAPSO), yielding a map and atomic model with resolution of 4.1 Å (EMDB 21494; PDB 6VZJ; Table S1; Fig. S2).

#### Cryo-EM structure determination: data collection and data reduction (NusG-TTC-A and NusG-TTC-B; n = 8; with CHAPSO)

Cryo-EM data for NusG-TTC-A and NusG-TTC-B (n = 8; with CHAPSO) were collected at the Rutgers University Cryo-EM core Facility, using a 200 kV Talos Arctica (FEI/ThermoFisher) electron microscope equipped with a GIF Quantum K2 direct electron detector (Gatan). Data collection was performed as in the preceding section. A total of 654 micrographs were recorded from one grid over one day. Micrographs were gain-normalized and defect-corrected.

Data were processed as summarized in Figs. S3A-D. Data processing was performed using a Tensor TS4 Linux GPU workstation with four GTX 1080 Ti graphic cards (NVIDIA). Dose weighting motion correction (3×3 tiles; b-factor = 150) were performed using Motioncor2 (*37*). CTF estimation was performed using CTFFIND-4.1 (*38*). Subsequent image processing was performed using Relion 3.0 (*39*). Automatic particle picking with Laplacian-of-Gaussian filtering yielded an initial set of 62,192 particles. Particles were extracted into 512×512 pixel boxes and subjected to rounds of reference-free 2D classification and removal of poorly populated classes, yielding a selected set of 10,874 particles. The selected set was 3D-classified with C1 symmetry, using a *de novo* 3D template created using 3D_initial_model under Relion 3.0. Classes that exhibited strong, well-defined density for the ribosome were selected. 9,278 particles for these classes were combined and 3D auto-refined using a mask with diameter of 450 Å, yielding a reconstruction with a global resolution that reached 5.8 Å, as determined from gold-standard Fourier shell correlation. The reconstruction showed clear density for the ribosome 70S subunit and less clear density for the TEC. The volume eraser tool of UCSF Chimera (*44*) was used to remove density for the 70S subunit, and a mask corresponding to the TEC was generated using relion_mask_create in Relion 3.0 with parameters --extend_inimask 8 and --width_soft_edge 3. The mask and the auto-refine optimizer star file were then employed for particle subtraction using relion_particle_subtraction in Relion 3.0, and the subtracted particles were used for masked 3D classification without image alignment. Two of five classes from masked 3D classification showed clear density for TEC and were selected separately and re-extracted. Several runs of auto-refinement, particle subtraction, and masked 3D classification were performed, yielding reconstructions at 6.3 Å (NusG-TTC-A) and 12.6 Å (NusG-TTC-B) (Fig. S3D; Table S1).

Initial atomic model building was performed as described in the preceding section, except that, for NusG-TTC-B, manual docking also was performed for NusG-C residues 123-181 from an NMR structure of *E. coli* NusB-S10 bound to NusG-C (PDB 2KVQ; *2*). Refinement of initial models was performed as in the preceding section, except that, for NusG-TTC-B, iterative cycles of model building and refinement also were performed for the NusG linker (residues 118-123) and NusG-C (residues 123-181).

The final atomic model of NusG-TTC-A (n = 8; with CHAPSO) was deposited in the EMDB and the PDB with accession codes EMDB 22193 and PDB 6XIJ (Table S1). The final atomic model of NusG-TTC-B (n = 8; with CHAPSO) was deposited in the EMDB and the PDB with accession codes EMDB 22192 and PDB 6XII (Table S1).

#### Cryo-EM structure determination: data collection and data reduction (NusG-TTC-B; n = 9 or 10; with CHAPSO)

Cryo-EM data for NusG-TTC-B (n = 9; with CHAPSO) were collected at the Rutgers University Cryo-EM core Facility, using a 200 kV Talos Arctica (FEI/ThermoFisher) electron microscope equipped with a GIF Quantum K2 direct electron detector (Gatan). Data collection was performed as in the preceding two sections, but with image collection in SerialEM (*48*) accelerated by use of coma-compensated beam-image shift (*49*). A total of 3,792 micrographs were recorded from one grid over three days. Micrographs were gain-normalized and defect-corrected.

Data were processed as summarized in Figs. S4A-D, using procedures as described in the preceding section. Data processing yielded a reconstruction at 4.7 Å resolution (Fig. S4D; Table S1).

Initial atomic model building and refinement of the initial model were performed as in the preceding section.

The final atomic model for NusG-TTC-B (n = 9; with CHAPSO) was deposited in the EMDB and the PDB with accession codes EMDB 22142 and PDB 6XDR (Table S1).

A cryo-EM structure of NusG-TTC-B (n = 10; with CHAPSO) was determined in the same manner as for NusG-TTC-B (n = 9; with CHAPSO), yielding a map and atomic model with resolution of 5.1 Å (EMDB 21469; PDB 6VYR; Table S1; Fig S4F).

#### Cryo-EM structure determination: data collection and data reduction (NusA-NusG-TTC-B; n = 8, 9, or 10; with CHAPSO)

Cryo-EM data for NusA-NusG-TTC-B (n = 8; with CHAPSO) were collected at NCCAT, using a 300 kV Krios Titan (FEI/ThermoFisher) electron microscope equipped with a Gatan K2 Summit direct electron detector (Gatan), Leginon (*47*), a nominal magnification of 105,000x, a calibrated pixel size of 0.4094 Å/pixel, and a dose rate of 45.44 electrons/ Å^2^/s. Movies were recorded at 30 ms/frame for 299 ms (50 frames), resulting in a total radiation dose of 68.15 electrons/Å^2^. Defocus range was varied between −1.25 µm and −2 µm. A total of 9,966 micrographs were recorded from one grid over three days. Micrographs were gain-normalized and defect-corrected.

Data were processed as summarized in Figs. S5A-D, using procedures as described in the preceding two sections. Data processing yielded reconstructions at 3.2 Å for NusA-NusG-TTC-B1, 3.5 Å for NusA-NusG-TTC-B2, and 3.1 Å for NusA-NusG-TTC-B3 (Fig. S5D; Table S1).

Initial atomic model building was performed as described in the preceding two sections, except that manual docking also was performed for RNAP α^I^ C-terminal domain (αCTD^I^), RNAP α^II^ C-terminal domains (αCTD^II^), and NusA segments from a cryo-EM structure of an *E. coli* paused TEC bound to NusA (PDB 6FLQ; *15*) and an NMR structure of αCTD bound to NusA AR2 (PDB 2JZB). Refinement of initial models was performed as in the preceding section, except that iterative cycles of model building and refinement also were performed for αCTD^I^, αCTD^II^, and NusA domains N (residues 1-136), S1, KH1, KH2, AR1, and AR2. Identities of αCTD^I^ and αCTD^I^ were assigned based on location; αCTD^I^ was assigned as the αCTD closest to residue 233 of α^I^ N-terminal domain, and αCTD^II^ was assigned as the αCTD closest to residue 233 of α^II^ N-terminal domain.

The final atomic models were deposited in the EMDB and PDB with accession codes EMDB 22082 and PDB 6X6T for NusA-NusG-TTC-B1 (n = 8; with CHAPSO), EMDB 22084 and PDB 6X7F for NusA-NusG-TTC-B2 (n = 8; with CHAPSO), and EMDB 22087 and PDB 6X7K for NusA-NusG-TTC-B3 (n = 9; with CHAPSO) (Table S1).

Cryo-EM structures for NusA-NusG-TTC-B (n = 10; with CHAPSO) were determined in the same manner, but using data collected at the Rutgers University Cryo-EM core Facility (data-collection procedures as in the preceding section; 4,736 micrographs from one grid over two days for n = 9; 2,624 micrographs from one grid over one day for n = 10), yielding maps and atomic models with resolutions of 5.9 Å for n = 9 NusG-TTC-B1, 4.2 Å for n = 9 NusG-TTC-B2, 4.8 Å for n = 9 NusG-TTC-B3 (EMDB 22107; PDB 6X9Q), 4.9 Å for n = 10 NusG-TTC-B1, and 3.7 Å for n = 10 NusG-TTC-B3 (EMDB 22141; PDB 6XDQ) (Table S1; Fig. S5G).

Cryo-EM data were collected and processed in the same manner for an analogous sample in the absence of NusG (n = 8; with CHAPSO), yielding a map and atomic model of TTC-A with resolution of 3.9 Å and a map and partial atomic model of NusA-TTC-X with resolution of 9.3 Å; Table S1; Fig. S8).

#### Cryo-EM structure determination: data collection and data reduction (NusG-TTC-A; n = 5 or 6; without CHAPSO)

Cryo-EM data for NusG-TTC-A (n = 5; without CHAPSO) were collected at the University of Michigan, Life Sciences Institute Cryo-EM Facility, using a 200 kV Glacios (FEI/ThermoFisher) electron microscope equipped with a K2 summit direct electron detector (Gatan). Data were collected using a stage tilt of 40°, in order to overcome particle orientation preferences that precluded structure determination using data collected without stage tilt (Fig. S12C). Following insertion of the grid into the microscope, eucentric height was obtained, and the stage was tilted to 40°C by changing the stage α parameter. Data were collected automatically in counting mode, using Leginon (*47*), a nominal magnification of 45,000x, a calibrated pixel size of 0.98 Å/pixel, and a dose rate of 7 electrons/^Å2^/s. Movies were recorded at 200 ms/frame for 6 s (30 frames total), resulting in a total radiation dose of 42 electrons/Å^2^. Defocus range was varied between −1 µm and −2 µm. A total of 1,212 micrographs were recorded from one grid over two days. Micrographs were gain-normalized and defect-corrected.

Data were processed as summarized in Figs. S12A-E. Dose weighting, motion correction (5×5 tiles; b-factor = 150) were performed using Motioncor2 (*37*). Per-particle CTF estimation was performed using goCTF (*50*), in order to estimate and account for the focus gradient in each tilted micrograph. Subsequent image processing was performed using Relion 3.0 (*39*). Automatic particle picking with Laplacian-of-Gaussian filtering yielded an initial set of 93,446 particles. Particles were binned 3x, extracted into 192−192 pixel boxes, and subjected to two rounds of reference-free 2D classification and removal of poorly populated classes, yielding a selected set of 41,868 particles. An *ab initio* model was generated using the SGD-based approach in Relion 3.0. 3D auto-refinement was performed against the selected set of particles, yielding a reconstruction with a global resolution that reached Nyquist frequency at 5.9 Å as determined from gold-standard Fourier shell correlation. Reference-free 2D classification and 3D auto-refinement also were performed with un-binned particles; this yielded a higher-resolution reconstruction, but did not yield clear improvement in subclass map quality. Therefore, subsequent steps were performed using subclass maps from 2D classification and 3D auto-refinement with binned data.

In the next step of data processing, particle subtraction was conducted to remove the signal from the ribosome 70S subunit, in order to characterize conformational states focused on the TEC. Because a mask accounting for the full range of conformational dynamics of the TEC was essential for the analysis, and because the segmented volume of the TEC was not reliable for generating such a mask (due to lower occupancy and higher conformational dynamics of the TEC as compared to the ribosome, and resulting weaker density for the TEC as compared to the ribosome in the map generated by 3D auto-refinement), a 3D volume was generated from signal-subtracted particles using the command relion_reconstruct, and this 3D volume was low-pass filtered to 60 Å resolution to generate a mask with an extended binary map in 9 pixels and a soft-edge in 3 pixels. The same 3D volume was low-pass filtered to 40 Å resolution and also used as an initial model for subsequent data processing. Focused 3D classifications without alignment was performed to sort different conformational states of the TEC relative to the ribosome. Of three subclasses obtained, two subclasses exhibited strong, well-defined density assignable to TEC: NusG-TTC-A1 (18%; identical to NusG-TTC-A of Figs. 1-2) and NusG-TTC-A2 (39%) (Fig. S12A). Non-signal-subtracted particles associated with each well-defined subclass were used to reconstruct a 3D volume using the command relion_reconstruct, and each volume was low-pass filtered to 30 Å resolution to generate an adaptive mask for subclass particles, with an extended binary map in 3 pixels and a soft-edge in 3 pixels. Local focused 3D refinement for each subclass then was performed with its adaptive mask using command relion_refine (-sigma_ang = 3). This process yielded reconstruction at 19 Å resolution for subclass A1 and at 14 Å resolution for subclass A2, as determined from gold-standard Fourier shell correlation (FSC; Fig. S12D; Table S2). Half maps were used to generate local-resolution maps using command relion_postprocess (-adhoc_bfac = 100) (Fig. S12E).

Initial atomic models for NusG-TTC-A1 and NusG-TTC-A2 (n = 5; without CHAPSO) were built by manual docking of RNAP β’, β, α^I^, α^II^ segments and NusG segments from a cryo-EM structure of an *E. coli* TEC bound to NusG (PDB 6C6U; *40*); DNA and RNA segments from a cryo-EM structure of an *E. coli* 21Q transcription antitermination “loaded” complex (PDB 6P19; *18*); ribosome 30S S2-S21 segments and 16S rRNA segments from a cryo-EM structure of an *E. coli* “expressome” (PDB 5MY1; *12*); ribosome 50S subunit from a cryo-EM structure of an *E. coli* 50S ribosomal subunit (PDB 6QDW); and P- and E-site tRNA segments from a crystal structure of a *Thermus thermophilus* 70S ribosome P- and E-site tRNA and mRNA (PDB 4V6G, *43*), using UCSF Chimera (*44*). For the NusG linker and NusG-C (residues 117-182), density was absent, suggesting high segmental flexibility; the NusG linker and NusG-C segments were not fitted. RNAP SI2 (residues 936-1041) was subjected to iterative cycles of model building and refinement in Coot (*46*). The final atomic model for TTC-A2 (n = 5) has been deposited in the EMDB and PDB with accession codes EMDB 21471 and PDB 6VYT (Table S2).

Cryo-EM structures for NusG-TTC-A (n = 6; without CHAPSO) were determined in the same manner, yielding maps and atomic models with resolutions of 15 Å for NusG-TTC-A1 and 13 Å for NusG-TTC-A2 (Table S2).

#### Cryo-EM structure determination: data collection and data reduction (NusG-TTC-A, NusG-TTC-C, and NusG-TTC-D; n =7; without CHAPSO)

Cryo-EM data for NusG-TTC-A, NusG-TTC-C and NusG-TTC-D (n = 7; without CHAPSO) were collected at the University of Michigan, Life Sciences Institute Cryo-EM Facility. Data collection using a stage tilt of 40° was performed as in the preceding section. A total of 7,297 micrographs were recorded from one grid over five days. Micrographs were gain-normalized and defect-corrected.

Data were processed as summarized in Figs. S13A-E. Dose weighting, motion correction (5×5 tiles; b-factor = 150) were performed using Motioncor2 (*37*). Per-particle CTF estimation was performed using goCTF (*50*), in order to estimate and account for the focus gradient in each tilted micrograph. Subsequent image processing was performed using Relion 3.0 (*39*). Automatic particle picking with Laplacian-of-Gaussian filtering yielded an initial set of 765,225 particles. Particles were binned 3x, extracted into 192×192 pixel boxes, and subjected to two rounds of reference-free 2D classification and removal of poorly populated classes, yielding a selected set of 445,738 particles. An *ab initio* model was generated using the SGD-based approach in Relion 3.0. 3D auto-refinement was performed against the selected set of particles, yielding a reconstruction with a global resolution that reached Nyquist frequency at 5.9 Å as determined from gold-standard Fourier shell correlation. Reference-free 2D classification and 3D auto-refinement also were performed with un-binned particles; this yielded a higher-resolution reconstruction, but did not yield clear improvement in subclass map quality. Therefore, subsequent steps were performed using subclass maps from 2D classification and 3D auto-refinement with binned data.

In the next step of data processing, particle subtraction was conducted to remove the signal from the ribosome 70S subunit, in order to characterize conformational states focused on the TEC. Because a mask accounting for the full range of conformational dynamics of the TEC was essential for the analysis, and because the segmented volume of the TEC was not reliable for generating such a mask (due to lower occupancy and higher conformational dynamics of the TEC as compared to the ribosome, and resulting weaker density for the TEC as compared to the ribosome in the map generated by 3D auto-refinement), a 3D volume was generated from signal-subtracted particles using the command relion_reconstruct, and this 3D volume was low-pass filtered to 60 Å resolution to generate a mask with an extended binary map in 9 pixels and a soft-edge in 3 pixels. The same 3D volume was low-pass filtered to 40 Å resolution and also used as an initial model for subsequent data processing. Focused 3D classifications without alignment was performed to sort different conformational states of the TEC relative to the ribosome. Of seven subclasses obtained, six subclasses exhibited strong, well-defined density assignable to the TEC: NusG-TTC-A2 (5%), NusG-TTC-C4 (2.9%), NusG-TTC-C5 (2.8%), NusG-TTC-C6 (2.7%), NusG-TTC-D2 (3.2%), and NusG-TTC-D3 (3.3%). The three NusG-TTC-C subclasses were grouped together and subjected to an additional round of 3D classification into three subclasses without alignment. All well-defined subclasses of RNAP were selected (Figs S13A), and non-signal-subtracted particles associated with each subclass were used to reconstruct a 3D volume using the command relion_reconstruct, each volume was low-pass filtered to 30 Å resolution to generate an adaptive mask for subclass particles, with an extended binary map in 3 pixels and a soft-edge in 3 pixels. Local focused 3D refinement for each subclass was then performed with its adaptive mask using command relion_refine (-sigma_ang = 3). This process yielded reconstructions at 11 Å resolution for subclass A2; at 10 Å resolution for subclasses C4, C5, and C6; and at 9 Å resolution for subclasses D2 and D3, as determined from gold-standard Fourier shell correlation (FSC; Figs S13D; Table S2). Half maps were used to generate local-resolution maps using command relion_postprocess (-adhoc_bfac = 100) (Fig. S13E).

Atomic models for NusG-TTC-A2, NusG-TTC-C4, NusG-TTC-C5, NusG-TTC-C6, NusG-TTC-D2, and NusG-TTC-D3 (n = 7; without CHAPSO) were built as described in the preceding section, yielding maps and atomic models with resolutions of 14 Å for NusG-TTC-A2, 9.9 Å for NusG-TTC-C4 (EMDB 21475; PDB 6VYX), 9.9 Å for NusG-TTC-C5 (EMDB 21476; PDB 6VYY), 9.9 Å for NusG-TTC-C6 (EMDB 21477; PDB 6VYZ), 8.9 Å for NusG-TTC-D2, and 8.9 Å for NusG-TTC-D3 (EMDB 21485; PDB 6VZ5) (Table S2).

#### Cryo-EM structure determination: data collection and data reduction (NusG-TTC-C, and NusG-TTC-D; n = 8 or 9; without CHAPSO)

Cryo-EM data for NusG-TTC-C and NusG-TTC-D (n = 9; without CHAPSO) were collected at the University of Michigan, Life Sciences Institute Cryo-EM Facility. Data collection using a stage tilt of 40° was performed as in the preceding two sections. A total of 11,989 micrographs were recorded from one grid over five days. Micrographs were gain-normalized and defect-corrected.

Data were processed as summarized in Figs. S14A-E. Dose weighting, motion correction (5×5 tiles; b-factor = 150) were performed using Motioncor2 (*37*). Per-particle CTF estimation was performed using goCTF (*50*), in order to estimate and account for the focus gradient in each tilted micrograph. Subsequent image processing was performed using Relion 3.0 (*39*). Automatic particle picking with Laplacian-of-Gaussian filtering yielded an initial set of 1,200,794 particles. Particles were binned 3x, extracted into 192×192 pixel boxes, and subjected to two rounds of reference-free 2D classification and removal of poorly populated classes, yielding a selected set of 569,067 articles. An *ab initio* model was generated using the SGD-based approach in Relion 3.0. 3D auto-refinement was performed against the selected set of particles, yielding a reconstruction with a global resolution that reached Nyquist frequency at 5.9 Å as determined from gold-standard Fourier shell correlation. Reference-free 2D classification and 3D auto-refinement also were performed with un-binned particles; this yielded a higher-resolution reconstruction, but did not yield clear improvement in subclass map quality. Therefore, subsequent steps were performed using subclass maps from 2D classification and 3D auto-refinement with binned data.

In the next step of data processing, particle subtraction was conducted to remove the signal from the ribosome 70S subunit, in order to characterize conformational states focused on the TEC. Because a mask accounting for the full range of conformational dynamics of the TEC was essential for the analysis, and because the segmented volume of the TEC was not reliable for generating such a mask (due to lower occupancy and higher conformational dynamics of the TEC as compared to the ribosome, and resulting weaker density for the TEC as compared to the ribosome in the map generated by 3D auto-refinement), a 3D volume was generated from signal-subtracted particles using the command relion_reconstruct, and this 3D volume was low-pass filtered to 60 Å resolution to generate a mask with an extended binary map in 9 pixels and a soft-edge in 3 pixels. The same 3D volume was low-pass filtered to 40 Å resolution and also used as an initial model for subsequent data processing. Focused 3D classifications without alignment was performed to sort different conformational states of the TEC relative to the ribosome. Of six subclasses obtained, five subclasses exhibited strong, well-defined density assignable to the TEC: NusG-TTC-C1 (2%), NusG-TTC-C2 (3%), NusG-TTC-C3 (6%), NusG-TTC-D1 (10%), and NusG-TTC-D2 (12%). The three NusG-TTC-C subclasses were grouped together and subjected to an additional round of 3D classification into three subclasses without alignment. All well-defined subclasses of RNAP were selected (Figs S14A), and non-signal-subtracted particles associated with each subclass were used to reconstruct a 3D volume using the command relion_reconstruct, each volume was low-pass filtered to 30 Å resolution to generate an adaptive mask for subclass particles, with an extended binary map in 3 pixels and a soft-edge in 3 pixels. Local focused 3D refinement for each subclass was then performed with its adaptive mask using command relion_refine (-sigma_ang = 3). This process yielded reconstructions at 7.6 Å resolution for subclass C1, at 7.0 Å resolution for subclasses C2 and C3, and at 10 Å resolution for subclasses D1 and D2, as determined from gold-standard Fourier shell correlation (FSC; Figs. S14D; Table S2). Half maps were used to generate local-resolution maps using command relion_postprocess (-adhoc_bfac = 100) (Fig. S14E).

Atomic models for NusG-TTC-C1, NusG-TTC-C2, NusG-TTC-C3, NusG-TTC-D1, and NusG-TTC-D2 (n = 9; without CHAPSO) were built as described in the preceding two sections. The final atomic models for NusG-TTC-C1, NusG-TTC-C2, NusG-TTC-C3, NusG-TTC-D1, and NusG-TTC-D2 (n = 9; without CHAPSO) have been deposited in the Electron Microscopy Data Bank (EMDB) and the PDB with accession codes EMD 21486 and PDB 6VZ7, EMD 21472 and PDB 6VYU, EMD 21474 and PDB6VYW, EMD-21482 and PDB 6VZ2, and EMD 21483 and PDB 6VZ3, respectively (Table S2).

Cryo-EM structures for NusG-TTC-C and NusG-TTC-D (n = 8; without CHAPSO) were determined in the same manner, yielding maps and atomic models with resolutions of 7.0 Å for NusG-TTC-C2, 7.0 Å for NusG-TTC-C3, 9.9 Å for NusG-TTC-C6, 10 Å for NusG-TTC-D2, 8.9 Å for NusG-TTC-D3, respectively (Table S2).

## Supplementary Figure Legends

**Fig. S1.**
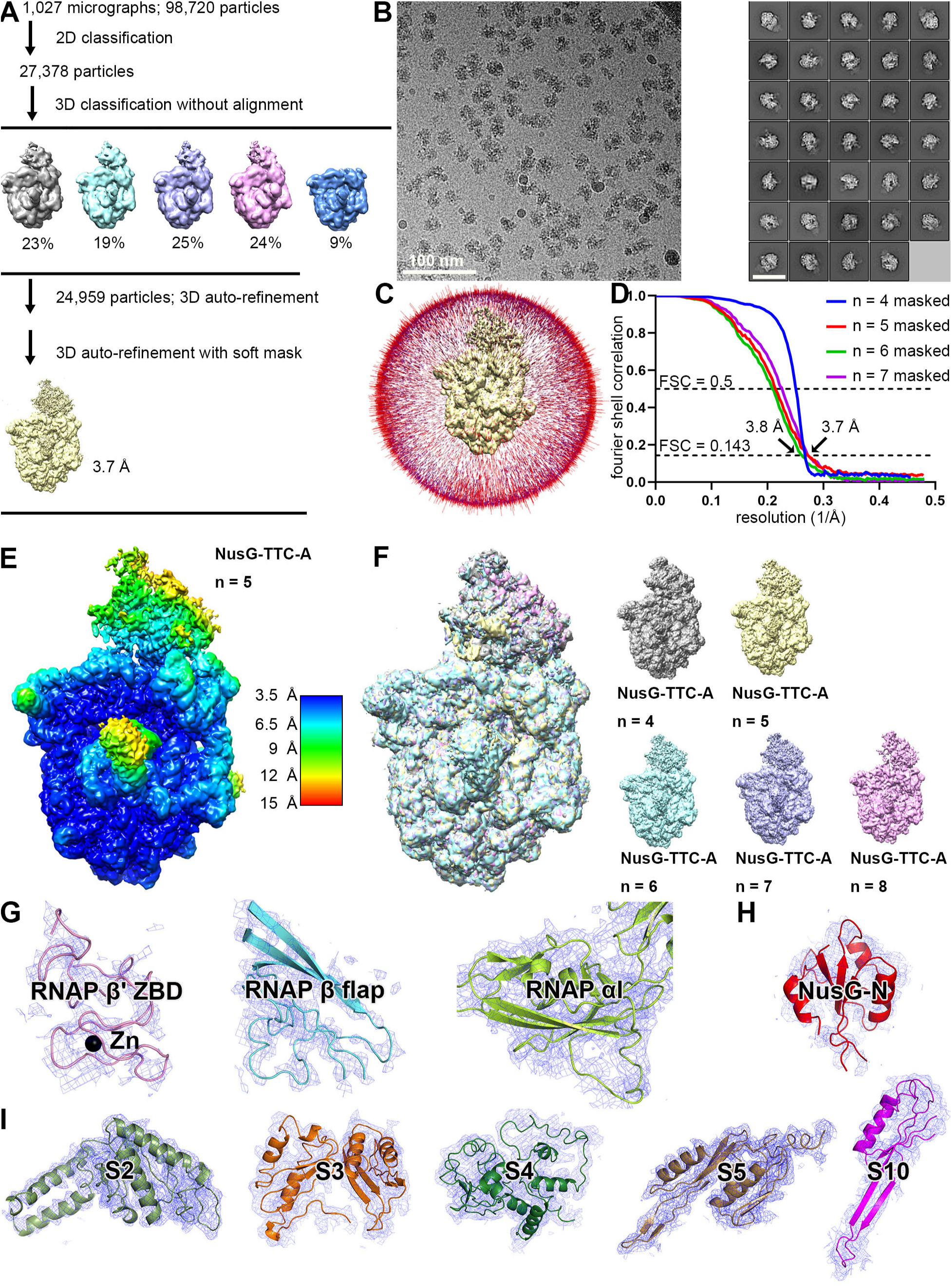
Structure determination: NusG-TTC-A (n = 4, 5, 6, or 7; with CHAPSO) **(A)** Data processing scheme (Table S1). **(B)** Representative electron micrograph and 2D class averages (50 nm scale bar in right subpanel). **(C)** Orientation distribution. **(D)** Fourier-shell-correlation (FSC) plot. **(E)** EM density map colored by local resolution. View orientation as in Figs. 1B and 2A, left. **(F)** EM density maps for NusG-TTC-A obtained using nucleic-acid scaffolds with n = 4, 5, 6, 7, and 8 (superimposition at left; individual EM maps and color scheme at right). View orientation as in Figs. 1A and 2A, left. **(G-I)** Representative EM density (blue mesh) and fits (ribbons) for RNAP regions that interact with ribosome, for NusG-N, and for ribosomal proteins that interact with RNAP. Data shown are for structure of NusG-TTC-A obtained with nucleic-acid scaffold having n = 5; similar data are obtained for structures of NusG-TTC-A obtained with nucleic-acid scaffolds having n = 4, 6, 7, or 8 (panels D and F; Fig. S3; Table S1).

**Fig. S2.**
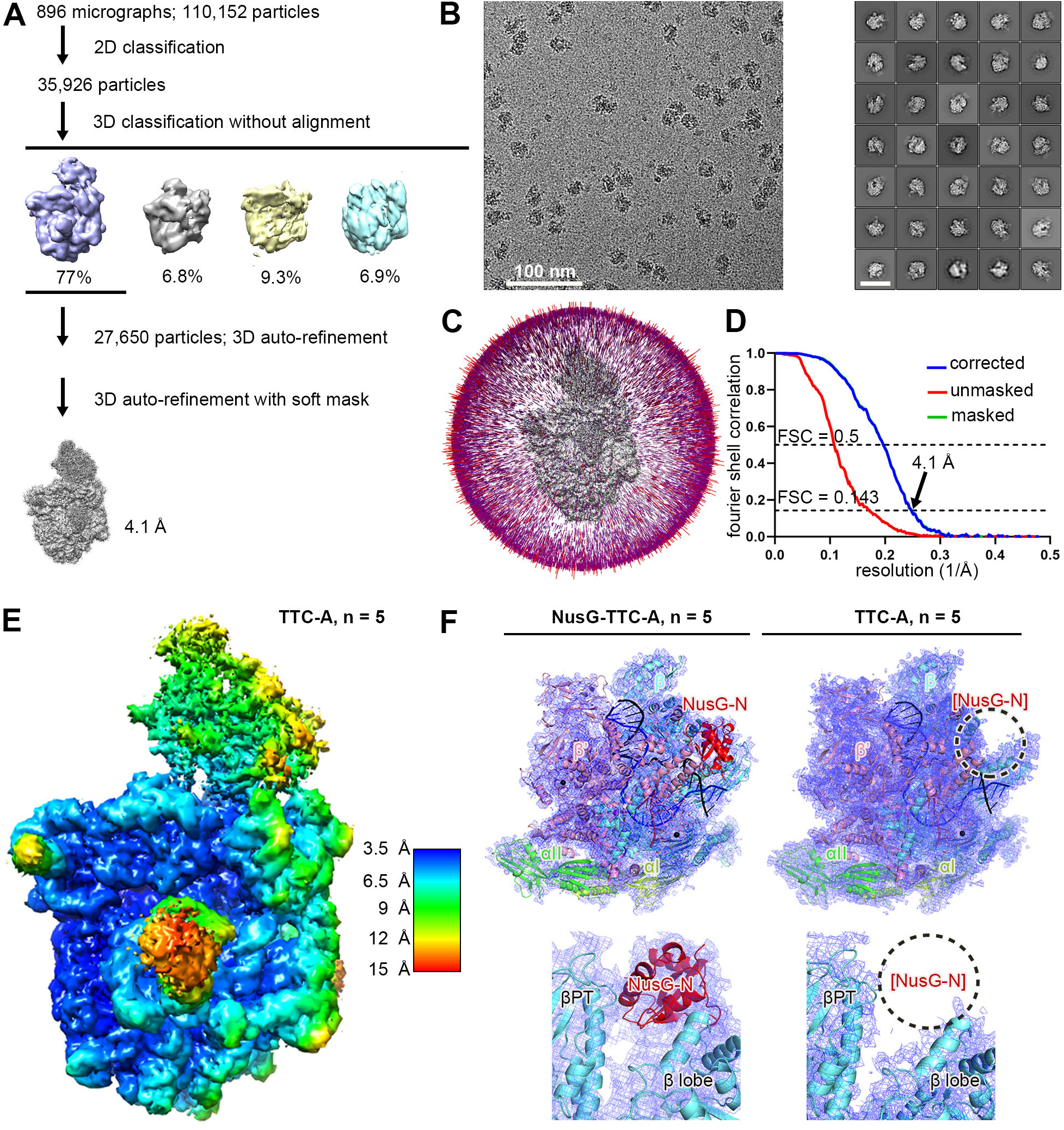
Structure determination: TTC-A in absence of NusG (n = 5; with CHAPSO) **(A)** Data processing scheme (Table S1). **(B)** Representative electron micrograph and 2D class averages (50 nm scale bar in right subpanel). **(C)** Orientation distribution. **(D)** Fourier-shell-correlation (FSC) plot. **(E)** EM density map colored by local resolution. View orientation as in Figs. 1B and 2A, left. **(F)** EM density and fit for NusG-TTC-A (left subpanel; overall view at top, close-up of NusG-N binding site below), and for TTC-A in absence of NusG (right; overall view at top, close-up of NusG-N binding site below). Blue mesh, EM density; dashed black circle, absent EM density for NusG-N in absence of NusG; βPT, RNAP β pincer tip. Other colors as in Figs. 1B and 2A.

**Fig. S3.**
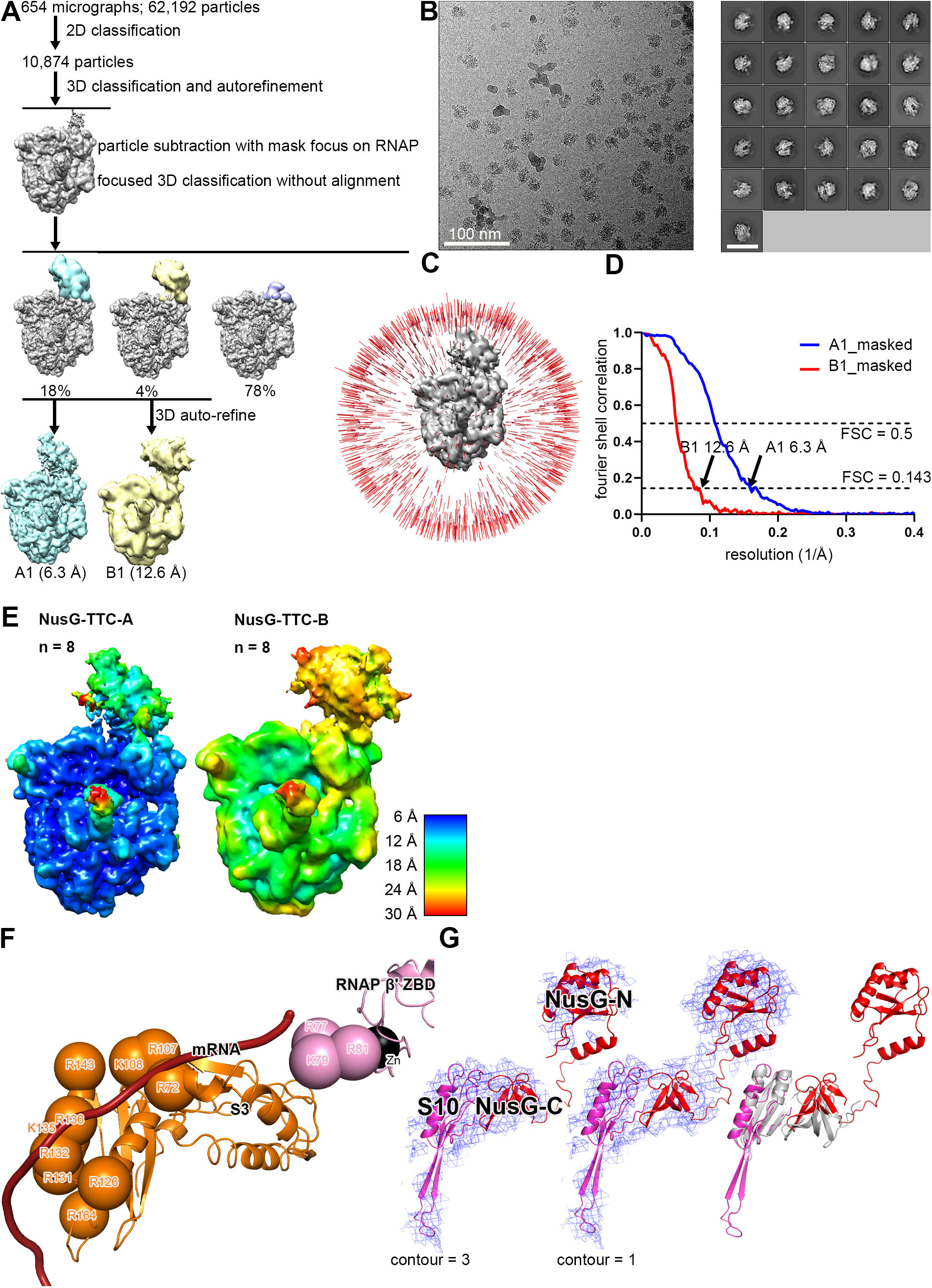
Structure determination: NusG-TTC-A and NusG-TTC-B (n = 8; with CHAPSO) **(A)** Data processing scheme (Table S1). Two classes were obtained: NusG-TTC-A (identical to NusG-TTC-A of Fig. S1) and NusG-TTC-B. **(B)** Representative electron micrograph and 2D class averages (50 nm scale bar in right subpanel). **(C)** Orientation distribution. **(D)** Fourier-shell-correlation (FSC) plot. **(E)** EM density maps colored by local resolution. View orientation as in Figs. 1B and 3A, left. **(F)** Path of mRNA (brick red) across RNAP-ribosome interface in NusG-TTC-B. Ribosomal protein S3, orange (positively charged residues positioned to contact mRNA as orange spheres); RNAP β’ zinc binding domain (ZBD, pink; Zn^2+^ ion as black sphere; positively charged residues positioned to contact mRNA as pink spheres). **(G)** Left, EM density (blue mesh) and fit for NusG (red) and ribosomal protein S10 (magenta) in NusG-TTC-B. Center, EM density at lower contour level and fit for NusG and ribosomal protein S10 in NusG-TTC-B (colored as in left subpanel). Right, superimposition of NusG and S10 of NusG-TTC-B (colored as in left subpanel) on NusG-C and S10 in NusG-C/S10 complex of *2* (PDB 2KVQ; gray).

**Fig. S4.**
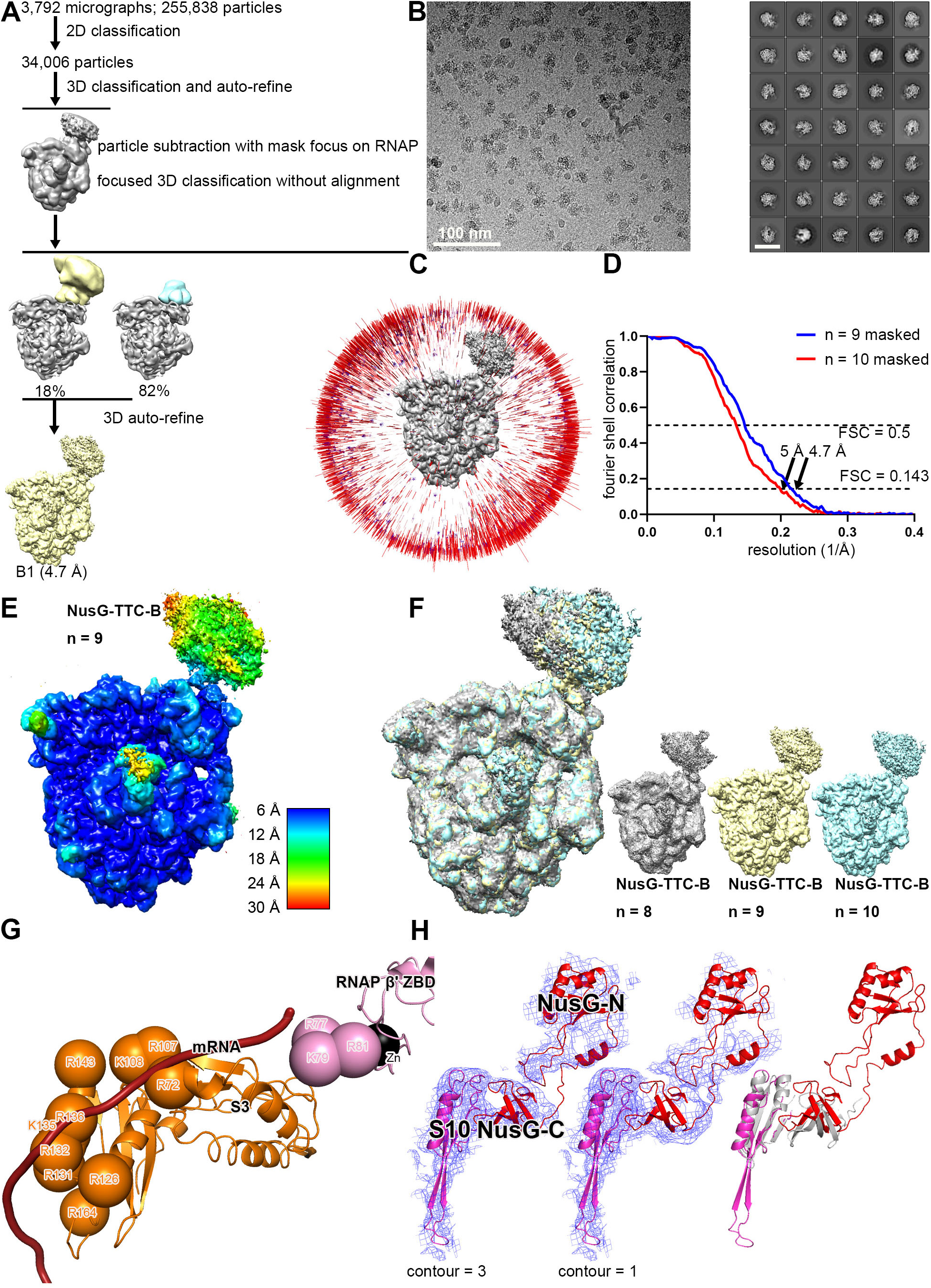
Structure determination: NusG-TTC-B (n = 9 or 10; with CHAPSO) **(A)** Data processing scheme (Table S1). **(B)** Representative electron micrograph and 2D class averages (50 nm scale bar in right subpanel). **(C)** Orientation distribution. **(D)** Fourier-shell-correlation (FSC) plot. **(E)** EM density map colored by local resolution. View orientation as in Figs. 1B and 3A, left. **(F)** EM density maps for NusG-TTC-B obtained using nucleic-acid scaffolds with n = 8, 9, or 10 (superimposition at left; individual EM maps and color scheme at right). View orientation as in Figs. 1A and 3A, left. **(G)** Path of mRNA (brick red) across RNAP-ribosome interface in NusG-TTC-B (n = 9). Ribosomal protein S3, orange (positively charged residues positioned to contact mRNA as orange spheres); RNAP β’ zinc binding domain (ZBD, pink; Zn^2+^ ion as black sphere; positively charged residues positioned to contact mRNA as pink spheres). **(H)** Left, EM density (blue mesh) and fit for NusG (red) and ribosomal protein S10 (magenta) in NusG-TTC-B (n = 9). Center, EM density at lower contour level and fit for NusG and ribosomal protein S10 in NusG-TTC-B (colored as in left subpanel). Right, superimposition of NusG and S10 of NusG-TTC-B (n = 9; colored as in left subpanel) on NusG-C and S10 of NusG-C/S10 complex of *2* (PDB 2KVQ; gray). Data shown are for structure of NusG-TTC-B obtained with nucleic-acid scaffold having n = 9; similar data are obtained with nucleic-acid scaffold having n = 10 (panels D and F; Table S1).

**Fig. S5.**
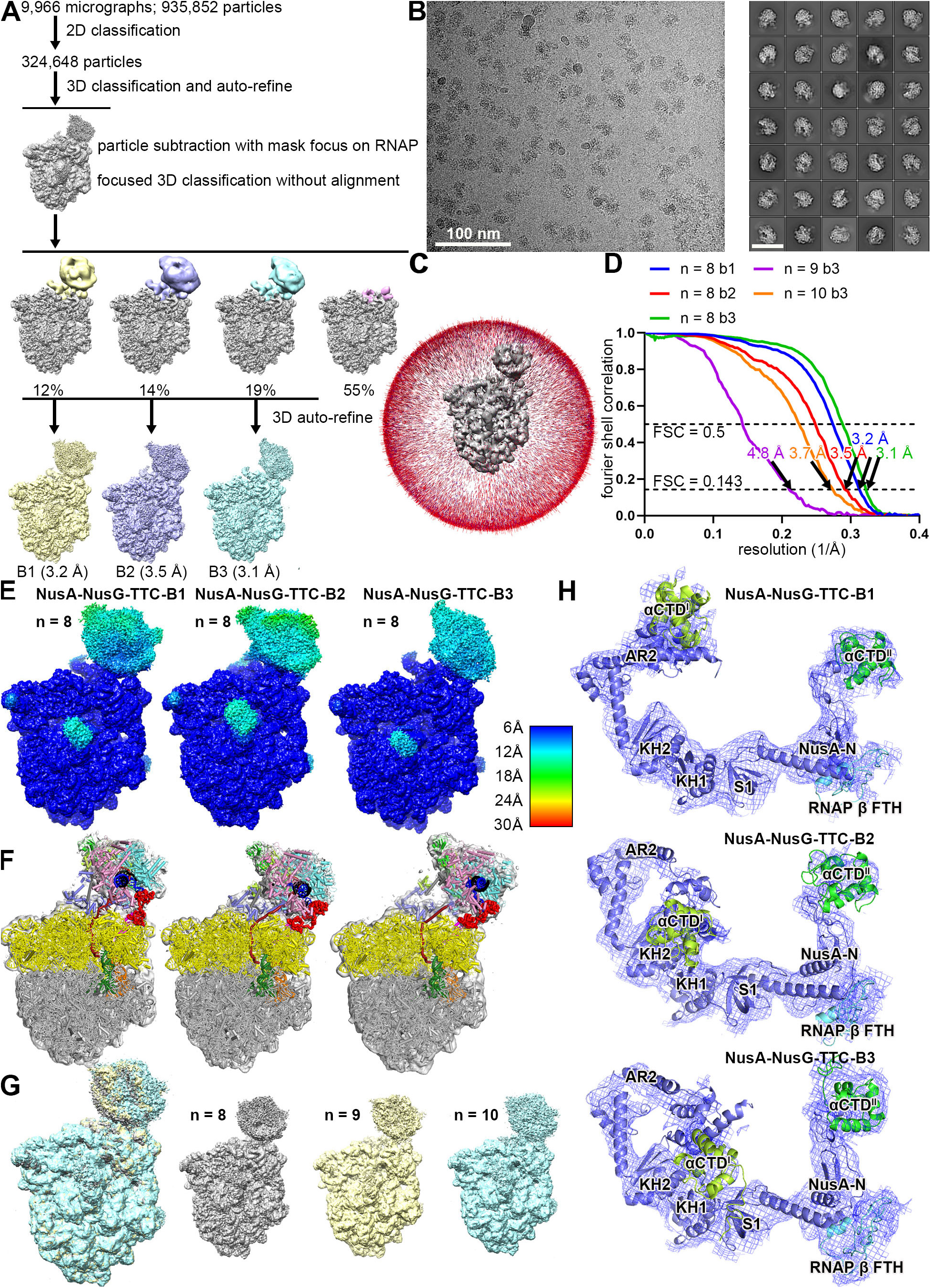
Structure determination: NusA-NusG-TTC-B (n = 8, 9, or 10; with CHAPSO) **(A)** Data processing scheme (Table S1). Three subclasses were obtained: NusA-NusG-TTC-B subclasses B1, B2, and B3. **(B)** Representative electron micrograph and 2D class averages (50 nm scale bar in right subpanel). **(C)** Orientation distribution. **(D)** Fourier-shell-correlation (FSC) plot. **(E)** EM density maps colored by local resolution. View orientation as in Figs. 1B and 4A, left. **(F)** EM density and fit for subclasses B1, B2, and B3. **(G)** EM density maps for NusA-NusG-TTC-B3 obtained using nucleic-acid scaffolds with n = 8, 9, or 10 (superimposition at left; individual EM maps and color scheme at right). View orientation as in Figs. 1B and 4A, left. **(H)** EM density and fit for NusA for subclasses B1, B2, and B3 (contour level 0.8 for EM density assigned to αCTD^I^ of subclass B1; contour level 3 for all other EM density). Data shown are for structures of TTC obtained with nucleic-acid scaffold having n = 8; similar data are obtained with nucleic-acid scaffold having n = 9 or 10 (panels D and G; Table S1).

**Fig. S6.**
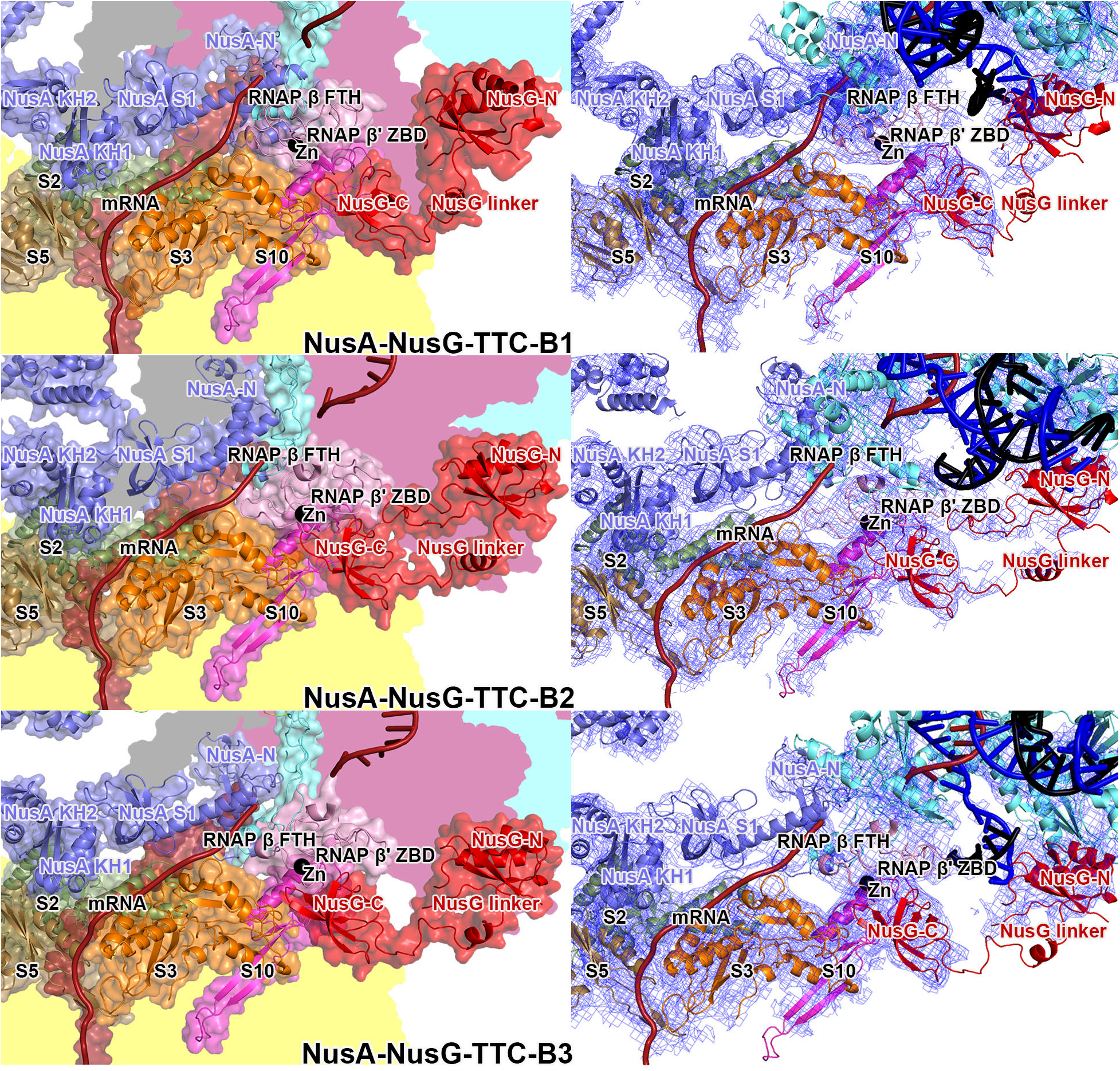
NusA-NusG-TTC-B subclasses B1, B2, and B3: RNAP-ribosome interface, NusG bridging, and NusA binding. RNAP-ribosome interface, NusG bridging, and NusA binding in NusA-NusG-TTC-B subclasses B1 (top), B2 (center), and B3 (bottom). Left subpanels, RNAP β’ zinc binding domain, (ZBD, pink; Zn^2+^ ion as black sphere) interacts with ribosomal protein S3 (orange) and mRNA (brick red). NusG (red) bridges RNAP and ribosome, with NusG-N interacting with RNAP and NusG-C interacting with ribosomal protein S10 (magenta). NusA (light blue) KH1 domain interacts with ribosomal proteins S5 and S2 (brown and forest green). Portions of RNAP β’, β, ω, and ribosome 30S not involved in interactions are shaded pink, cyan, gray, and yellow, respectively. Right subpanels, same, showing EM density as blue mesh.

**Fig. S7.**
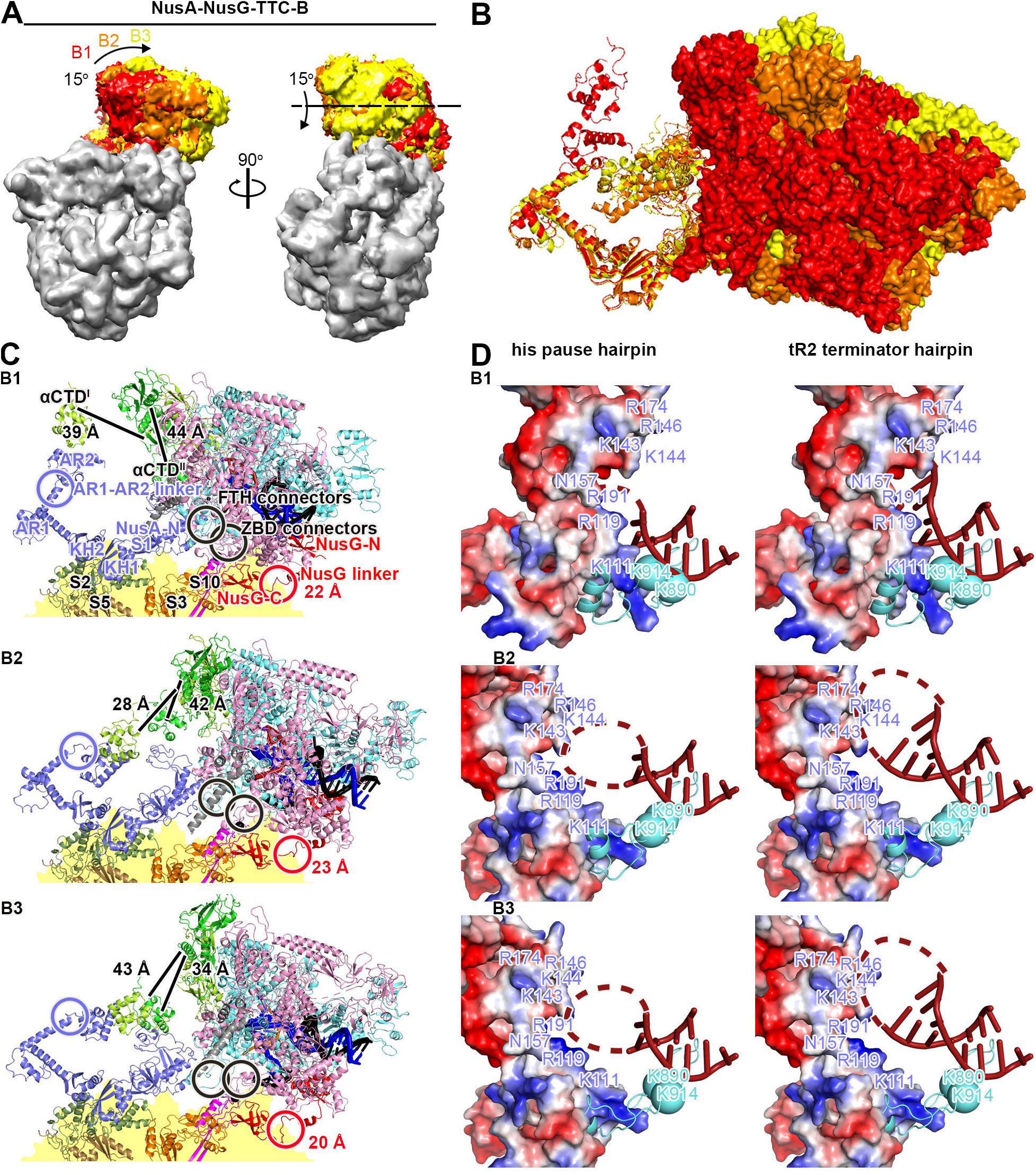
NusA-NusG-TTC-B subclasses B1, B2, and B3: structural relationship, points of flexibility, and accommodation of pause and terminator RNA hairpins. **(A)** Structural relationship of NusA-NusG-TTC-B subclasses B1, B2, and B3. Superimposition of EM maps of NusA-NusG-TTC-B subclasses B1, B2, and B3 (RNAP EM density colored by subclass; ribosome EM density in gray for subclass B2). Orientation of RNAP relative to ribosome in subclass B1 is most similar to that in NusG-TTC-B. View orientation as in Figs. 1B and 4A, left. See Movie S2. **(B)** Structural relationship of NusA-NusG-TTC-B subclasses B1, B2, and B3. Superimposition of NusA (ribbons) and RNAP (surfaces) of subclasses B1, B2, and B3. Colors as in (A). View orientation as in Fig. 4G. **(C)** Points of conformational difference--flexibility--in NusA-NusG-TTC-B subclasses B1 (top), B2 (center), and B3 (bottom): flexible linkage in NusA structure (AR1-AR2 linker; light blue circle), three flexible linkages between NusA and RNAP (β FTH connectors, αCTD^I^ linker, and αCTD^II^ linker; black circle and black lines), flexible linkage between RNAP and ribosome (β’ ZBD connectors; black circle), and flexible NusG bridging of RNAP and ribosome (NusG linker; red circle). View orientation as in Fig. 4G. **(D)** Modelled accommodation of *his* pause and tR2 terminator RNA hairpins (hairpin stem in red; hairpin loop as red dashed line) in NusA-NusG-TTC-B subclasses B1 (top; steric clash with *his* and tR2 harpins), B2 (center; no steric clash with *his* and tR2 harpins), and B3 (bottom; no steric clash with *his* and tR2 harpins) NusG, solvent-accessible surface colored by electrostatic potential (negative potential, red; positive potential, blue; positively charged residues, blue labels); RNAP β FTH, cyan ribbon (positively charged residues, cyan spheres and labels).

**Fig. S8.**
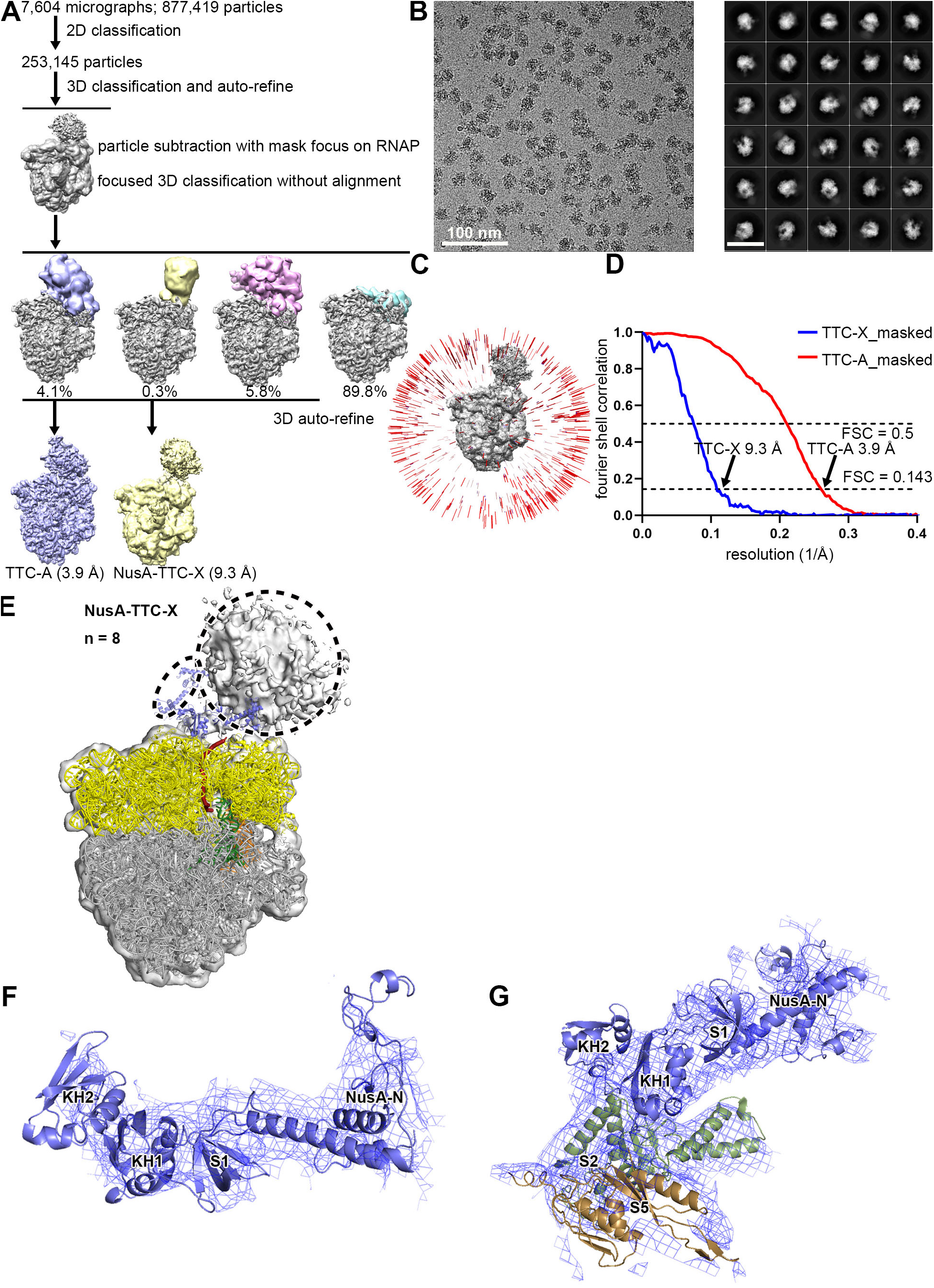
Structure determination: TTC-A and NusA-TTC-X (n = 8; with CHAPSO) **(A)** Data processing scheme (Table S1). Two classes were obtained: TTC-A (identical to TTC-A of Fig. S2) and NusA-TTC-X. **(B)** Representative electron micrograph and 2D class averages (50 nm scale bar in right subpanel). **(C)** Orientation distribution. **(D)** Fourier-shell-correlation (FSC) plot. **(E)** EM density and fit for NusA-TTC-X. NusA domains N, S1, KH1, and KH2, light blue; small dashed oval absent EM density for NusA domains AR1 and AR2; large dashed circle density assignable as, but not fittable as, RNAP. View orientation as in Figs. 1B and 4A, left. **(F)** EM density for NusA N, S1, KH1, and KH2 domains. View orientation as in Fig. S5H. **(G)** NusA-ribosome interface in NusA-TTC-X. NusA (light blue) KH1 domain interacts with ribosomal proteins S5 and S2 (brown and forest green). View orientation as in Fig. 4C.

**Fig. S9.**
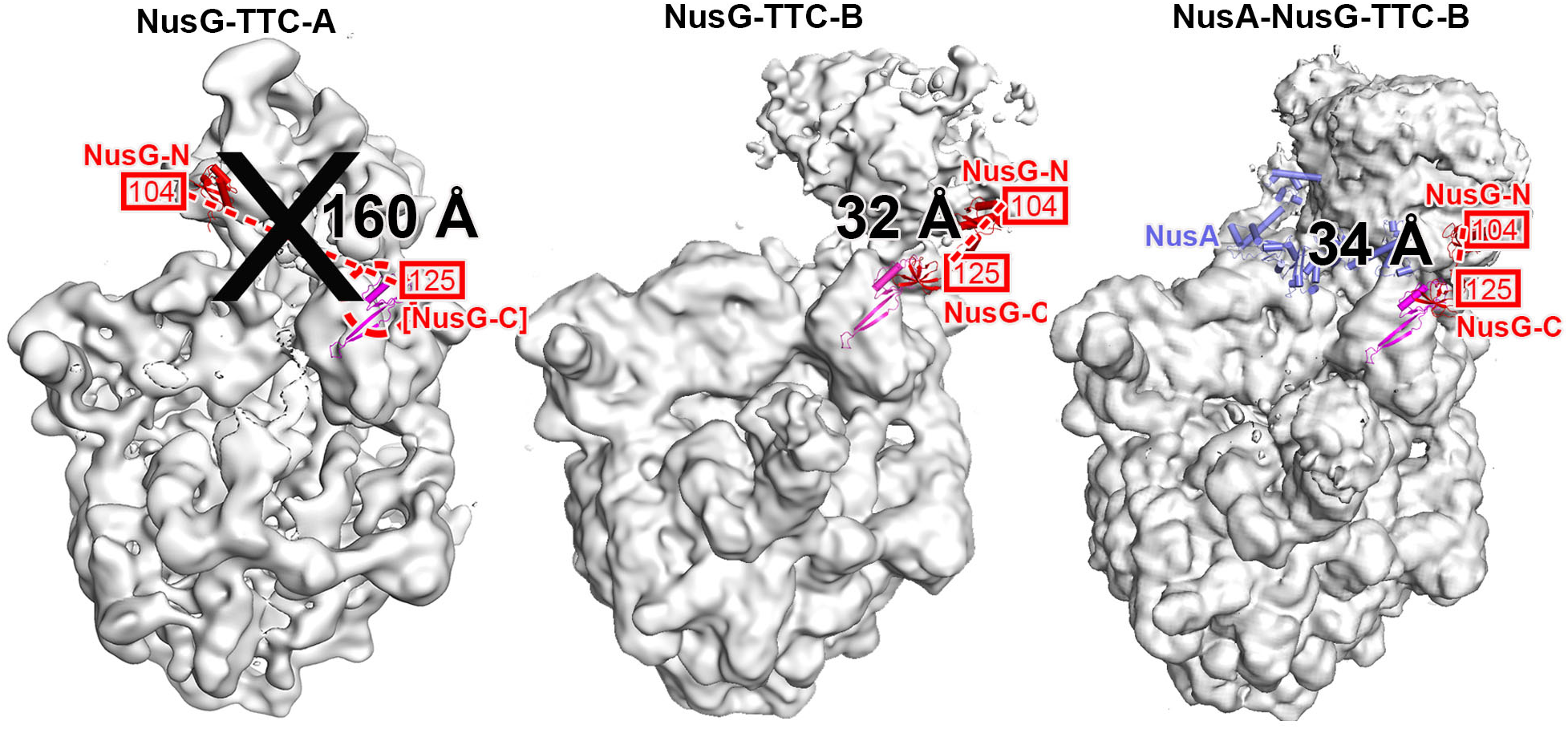
Compatibility of structures of NusG-TTC-A, NusG-TTC-B, and NusA-NusG-TTC-B with NusG bridging. Structures shown are NusG**-**TTC-A (subclass A1; left), NusG-TTC-B (center), and NusA-NusG-TTC-B (right). Black “X” indicates incompatibility with NusG bridging. NusG-N of NusG-TTC-A and NusG of NusG-TTC-B and NusA-NusG-TTC-B, red; ribosomal protein S10 (molecular target of NusG-C), magenta; modeled NusG-C in NusG-TTC-A, red with red dashed circle; measured length of shortest sterically allowed connection between NusG-N residue 104 and NusG-C residue 123, red dashed line and black distance in Å. The 22-residue linker between NusG-N residue 104 and NusG-C residue 125 can span 86 Å (22 residues x 3.9 Å/residue = 86 Å). The distance in NusG-TTC-A is incompatible with NusG bridging. The distances in NusG-TTC-B and NusA-NusG-TTC-B are compatible with NusG bridging.

**Fig. S10.**
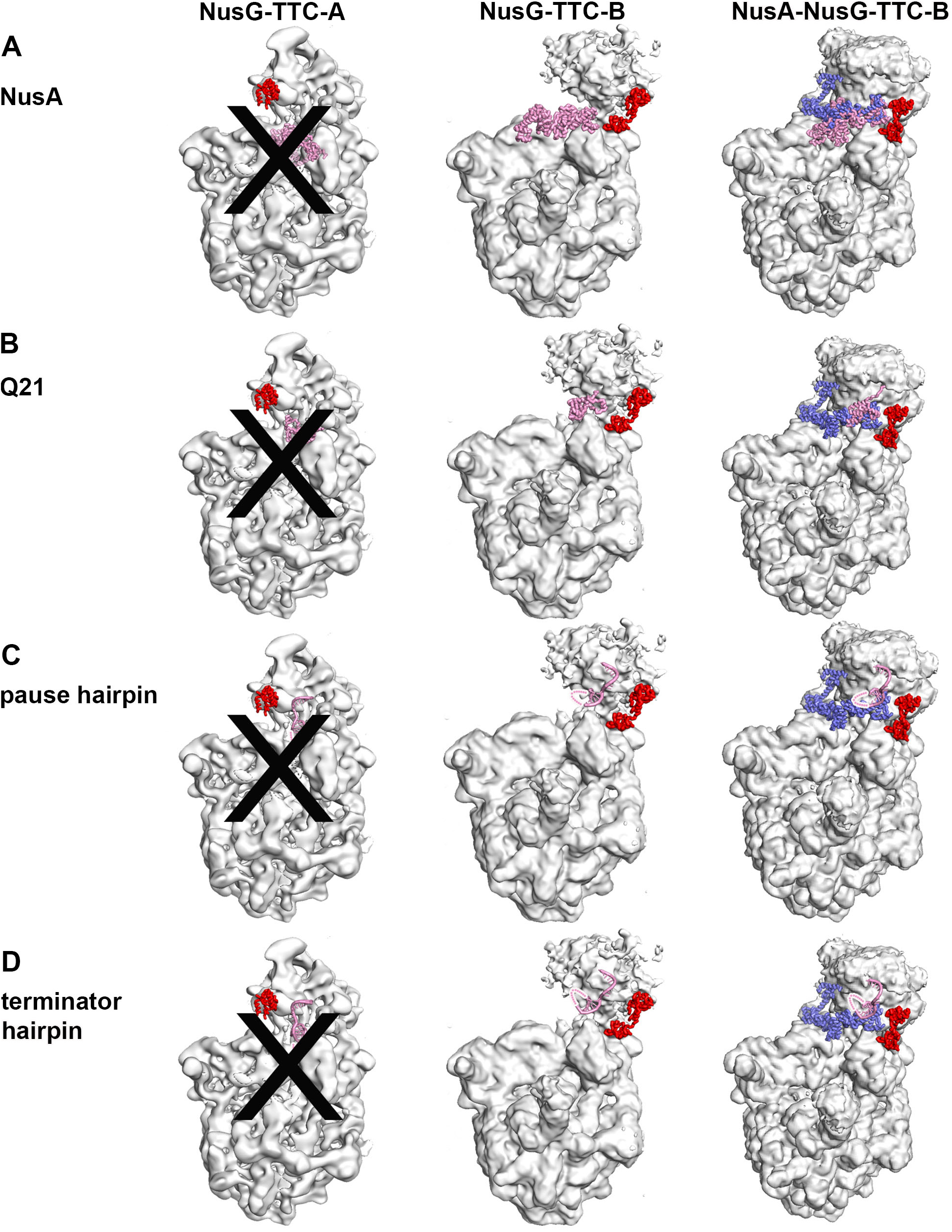
Compatibility of structures of NusG-TTC-A, NusG-TTC-B, and NusA-NusG-TTC-B with NusA binding formation of 21 Q antitermination complex, and formation of pause and termination RNA hairpins. Structures shown are NusG-TTC-A (subclass A1; left), NusG-TTC-B (center), and NusA-NusG-TTC-B (right). Black “X” indicates incompatibility. NusG-N of NusG-TTC-A and NusG of NusG-TTC-B and NusA-NusG-TTC-B, red; NusA of NusA-NusG-TTC-B, light blue; modelled NusA, 21 Q antitermination complex, or RNA hairpin, pink. **(A)** Compatibility with binding of transcription-elongation factor NusA (pink; *15*). NusG-TTC-A is incompatible with binding of NusA (severe steric clash of NusA N, S1, KH1, KH2, and AR1 with ribosome in subclass A1; severe steric clash of NusA KH2 and AR1 with ribosome in subclass A2). NusG-TTC-B (with very modest rotation of NusA) and, by definition, NusA-NusG-TTC-C are compatible with binding of NusA. **(B)** Compatibility with formation of bacteriophage 21 Q transcription antitermination complex (21 Q loaded complex; pink; *17*). NusG-TTC-A is incompatible with formation of 21 Q transcription antitermination complex (severe steric clash of Q body with ribosome in subclass A1 and subclass A2). NusG-TTC-B and NusA-NusG-TTC-B are compatible with formation of 21 Q transcription antitermination complex. **(C-D)** Compatibility with formation of His pause and tR2 terminator RNA hairpin (hairpin stem, pink; hairpin loop, pink dotted line; *15, 18–19*). NusG-TTC-A is incompatible with formation of pause and termination hairpins (collision of hairpin loop with ribosome in subclass A1). NusG-TTC-B and NusA-NusG-TTC-B are compatible with formation of pause and termination hairpins.

**Fig. S11.**
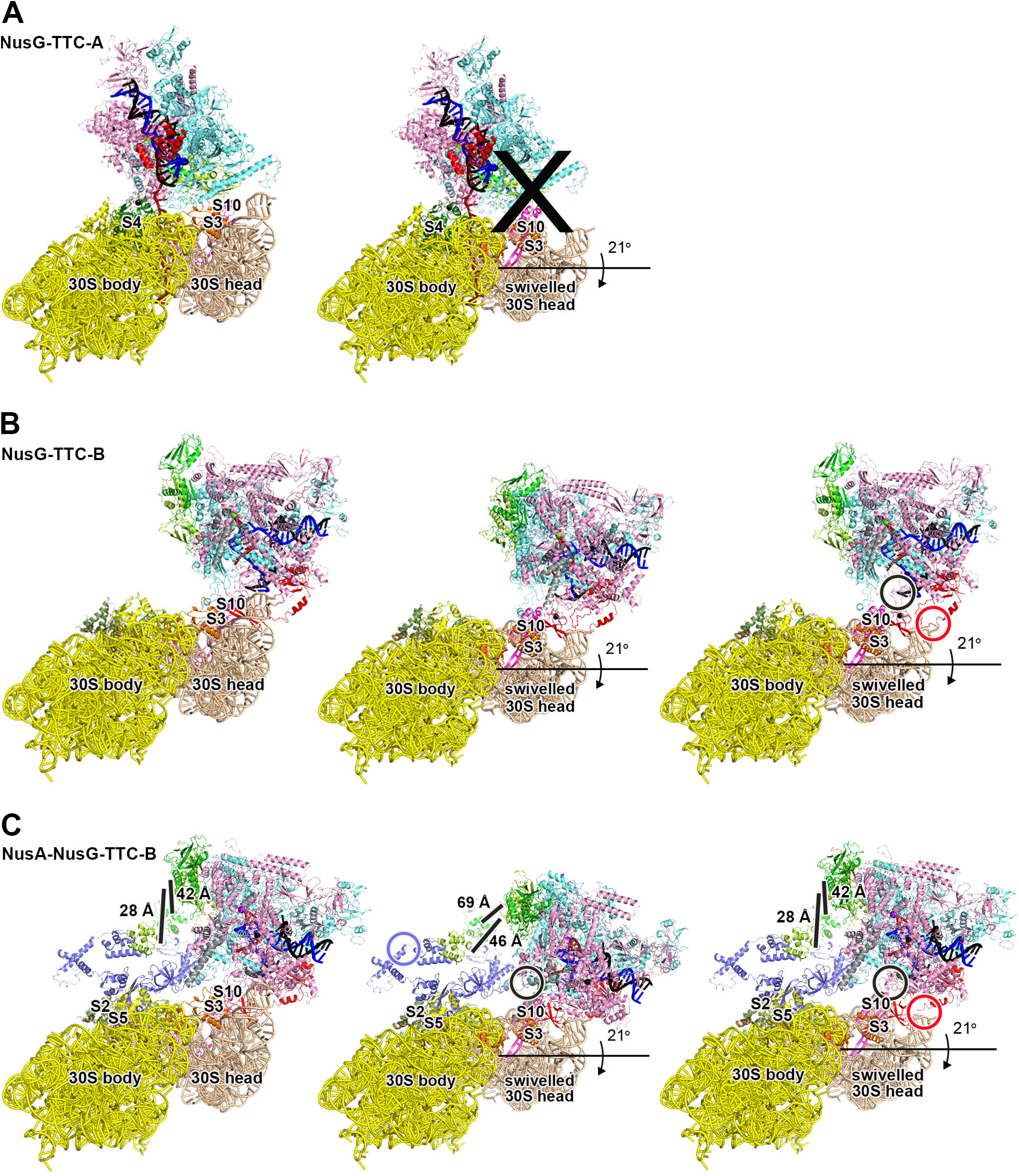
Compatibility of structures of NusG-TTC-A, NusG-TTC-B, and NusA-NusG-TTC-B with 30S-head swivelling. **(A)** NusG-TTC-A is incompatible with 30S-head swivelling, the ∼21° rotation of ribosome 30S head (brown) relative to ribosome 30S body (yellow) that occurs during ribosome translocation (severe disruption of RNAP-ribosome interactions). Structures of unswivelled and swivelled states are from PDB 4V51 and PDB 4W29, respectively (*51–52*). Ribosome 50S subunit and tRNAs are omitted for clarity. See Movie S3. **(B)** NusG-TTC-B is compatible with 30S-head swivelling (complete retention of RNAP-ribosome and NusG-ribosome interactions, provided RNAP and NusG move with 30S head (center subpanel), or provided RNAP β’ ZBD and NusG-C move with 30S head, exploiting flexible connectors between β’ ZBD and rest of RNAP and between NusG-N and NusG-C (right subpanel; black and red circles). See Movie S4. **(C)** NusA-NusG-TTC-B is compatible with 30S-head swivelling (complete retention of RNAP-ribosome, NusG-ribosome, and NusA-ribosome interactions, provided RNAP and NusG move with 30S head, exploiting flexible connectors within NusA and between NusA and RNAP (NusA “coupling pantograph”; center subpanel; light blue circle, black circle, and black lines), or provided RNAP β’ ZBD and NusG-C move with 30S head, exploiting flexible connectors between β’ ZBD and rest of RNAP and between NusG-N and NusG-C (right subpanel; black and red circles). See Movie S5.

**Fig. S12.**
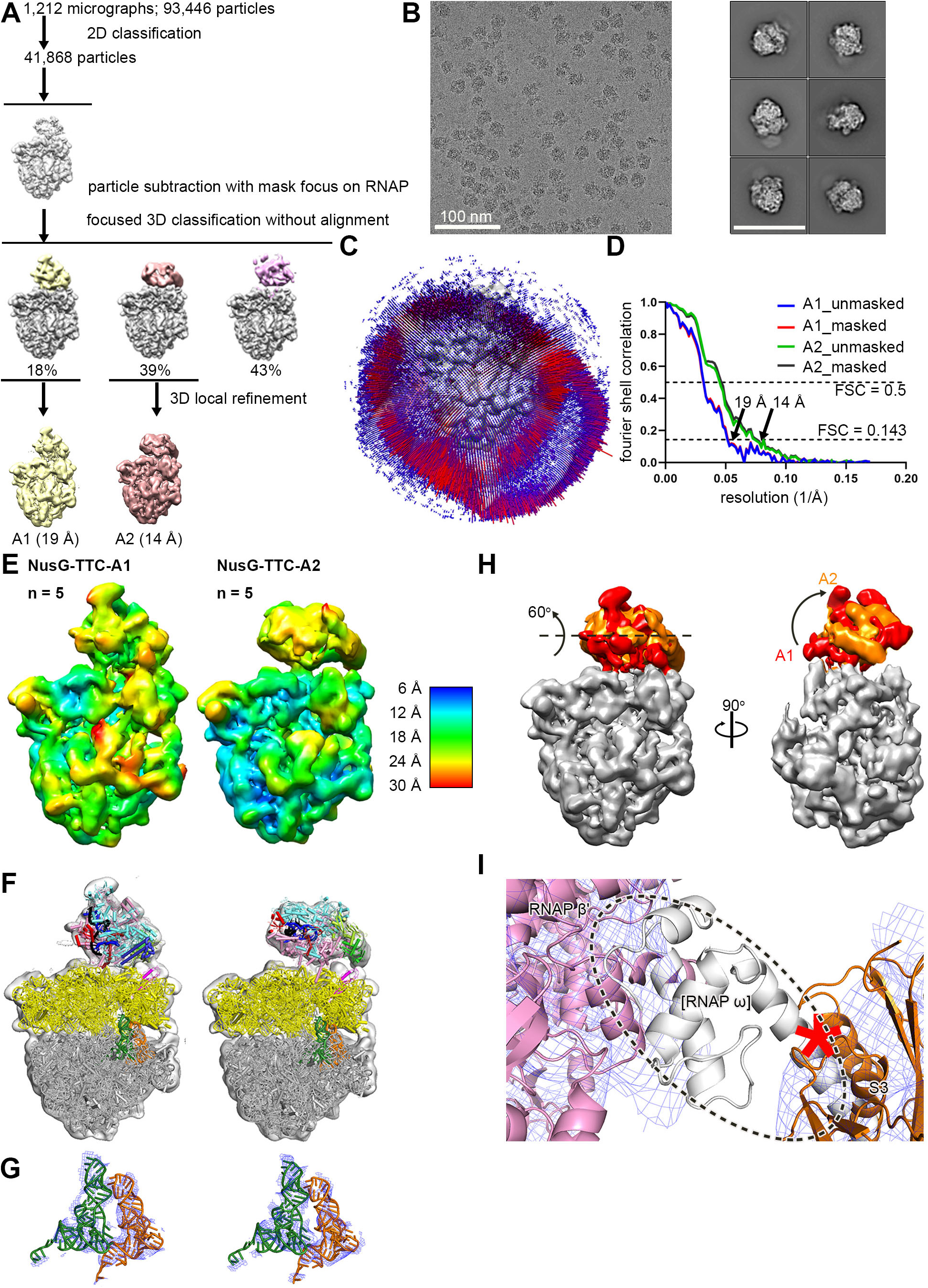
Structure determination: NusG-TTC-A (n = 5 or 6; without CHAPSO) **(A)** Data processing scheme (Table S2). Two subclasses were obtained: A1 and A2. **(B)** Representative electron micrograph and 2D class averages (50 nm scale bar in right subpanel). **(C)** Orientation distribution. **(D)** Fourier-shell-correlation (FSC) plot. **(E)** EM density maps colored by local resolution for subclasses A1 and A2. View orientation as in Figs. 1B and 2A, left. **(F)** EM density and fit for subclasses A1 and A2. **(G)** EM density (blue mesh) and fit (green, P-site tRNA; orange, E-site tRNA) for subclasses A1 and A2. **(H)** Superimposition of EM density maps of subclasses A1 (red) and A2 (orange). View orientations as in (A). **(I)** Absence of EM density for RNAP ω subunit in subclass A2. EM density, blue mesh; atomic models for RNAP β’ and ribosomal protein S2, pink ribbon and red-pink ribbon, respectively; location of missing EM for ω, dashed oval; ω in TEC in absence of ribosome (PDB 6P19; *19*), white ribbon. Analogous data for subclass A1 are in Fig 2E. Data shown are for structures of TTC obtained with nucleic-acid scaffold having n = 5; similar data are obtained for structures with nucleic-acid scaffold having n = 6 (Table S2).

**Fig. S13.**
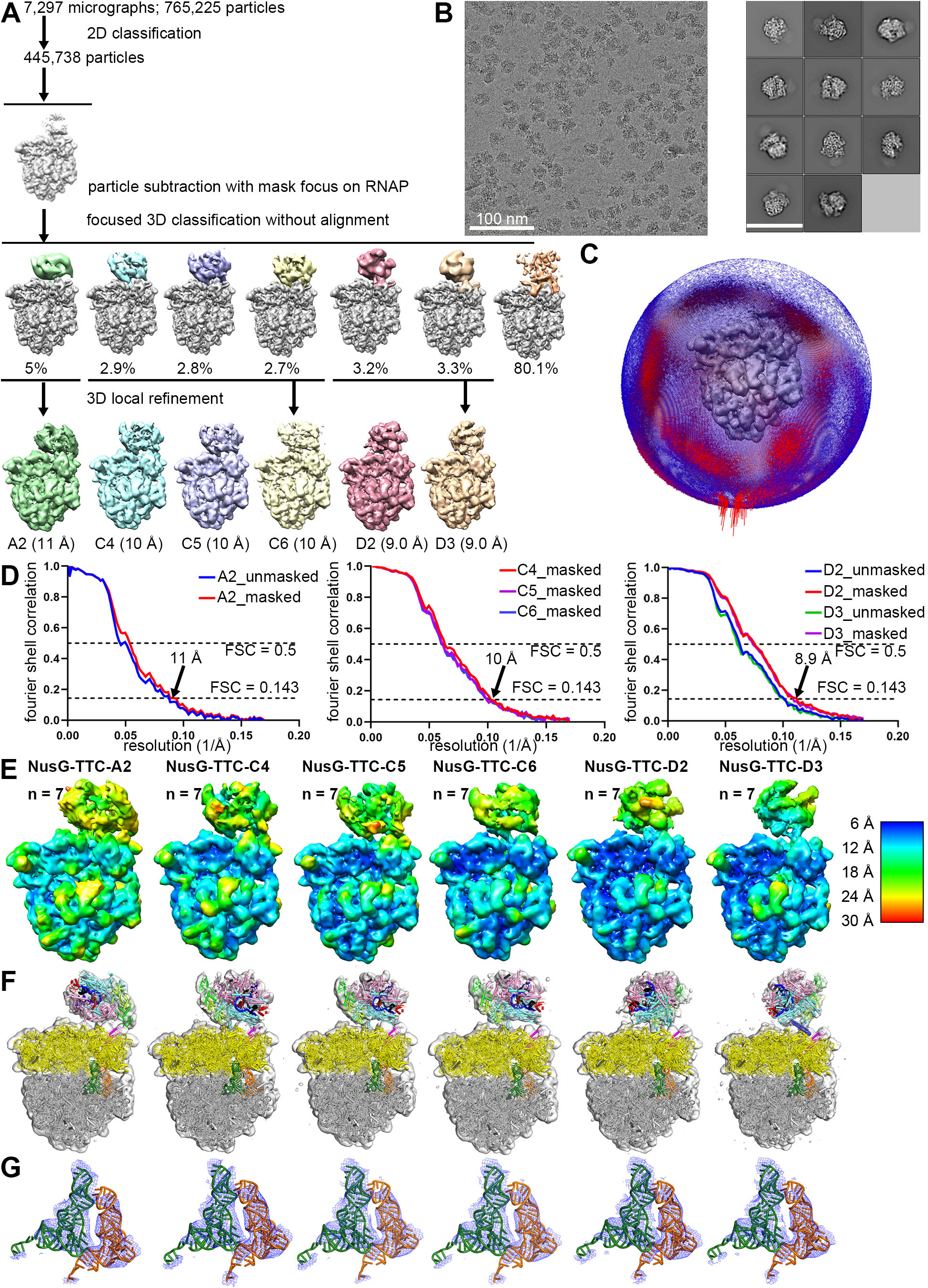
Structure determination: NusG-TTC-A, NusG-TTC-C, and NusG-TTC-D (n = 7; without CHAPSO) **(A)** Data processing scheme (Table S2). Six subclasses were obtained: NusG-TTC-A subclass A2; NusG-TTC-C subclasses C4, C5, and C6; and NusG-TCC-D subclasses D2 and D3. **(B)** Representative electron micrograph and 2D class averages (50 nm scale bar in right subpanel). **(C)** Orientation distribution. **(D)** Fourier-shell-correlation (FSC) plot. **(E)** EM density maps colored by local resolution. View orientation as in Figs. 1B, 2A, 3A, and 4A. **(F)** EM density and fit for subclasses A2, C4, C5, C6, D2, and D3. **(G)** EM density (blue mesh) and fit (green, P-site tRNA; orange, E-site tRNA) for subclasses A2, C4, C5, C6, D2, and D3.

**Fig. S14.**
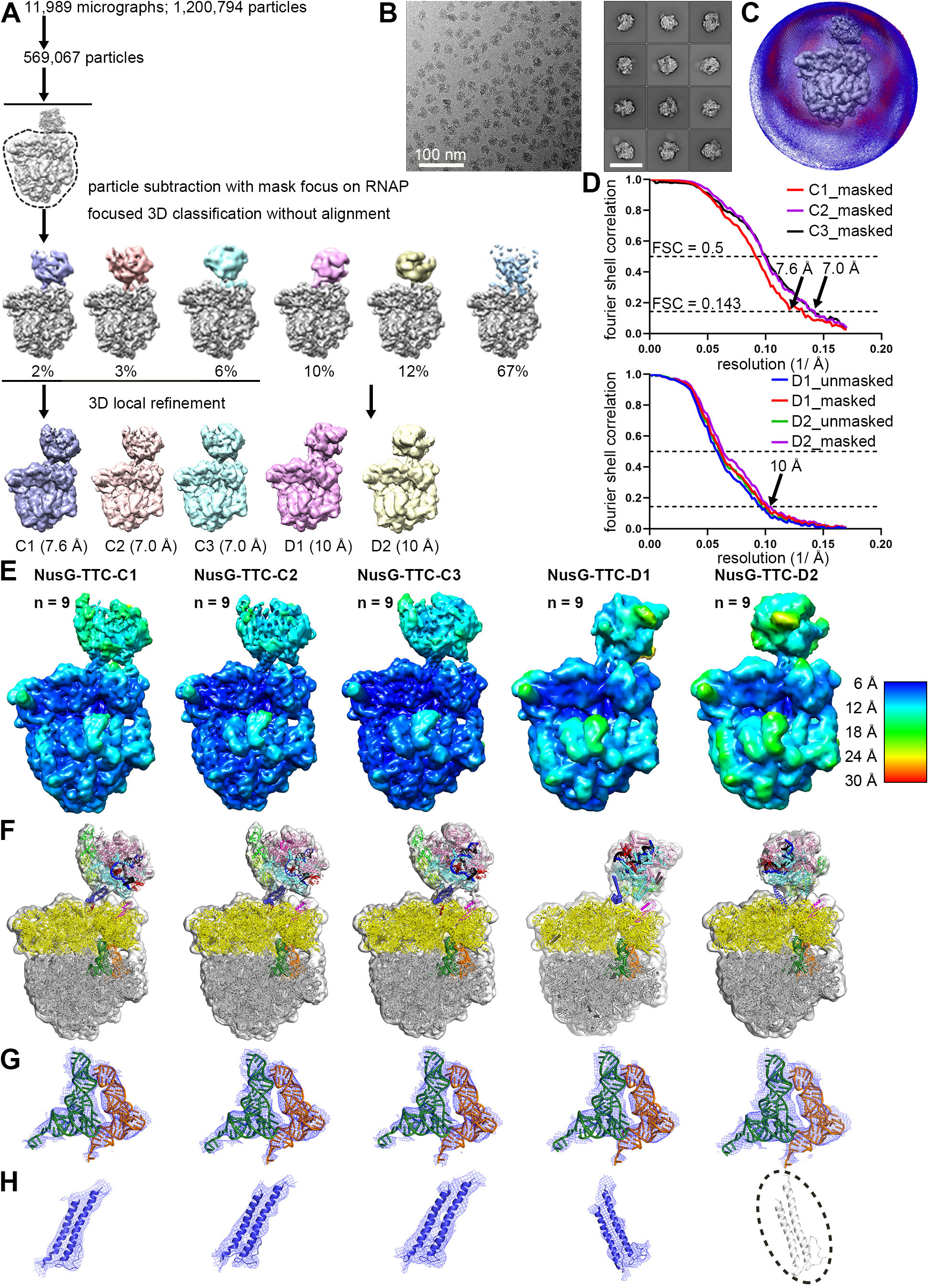
Structure determination: NusG-TTC-C and NusG-TTC-D (n = 8 or 9; without CHAPSO) **(A)** Data processing scheme (Table S2). Five subclasses were obtained: NusG-TTC-C subclasses C1, C2, and C3 and NusG-TTC-D subclasses D1 and D2. **(B)** Representative electron micrograph and 2D class averages (50 nm scale bar in right subpanel). **(C)** Orientation distribution. **(D)** Fourier-shell-correlation (FSC) plot. **(E)** EM density maps colored by local resolution. View orientation as in Figs. 1B, 3A, and 4A. **(F)** EM density and fit for subclasses C1, C2, C3, D1, and D2. **(G)** EM density (blue mesh) and fit (green and orange for P-site tRNA and E-site tRNA) for subclasses C1, C2, C3, D1, and D2. **(H)** EM density for βSI2 for subclasses C1, C2, C3, D1, and D2 (disordered in D2). Data shown are for structures of TTC obtained with nucleic-acid scaffold having n = 9; similar data are obtained with nucleic-acid scaffold having n = 8 (Table S2).

**Fig. S15.**
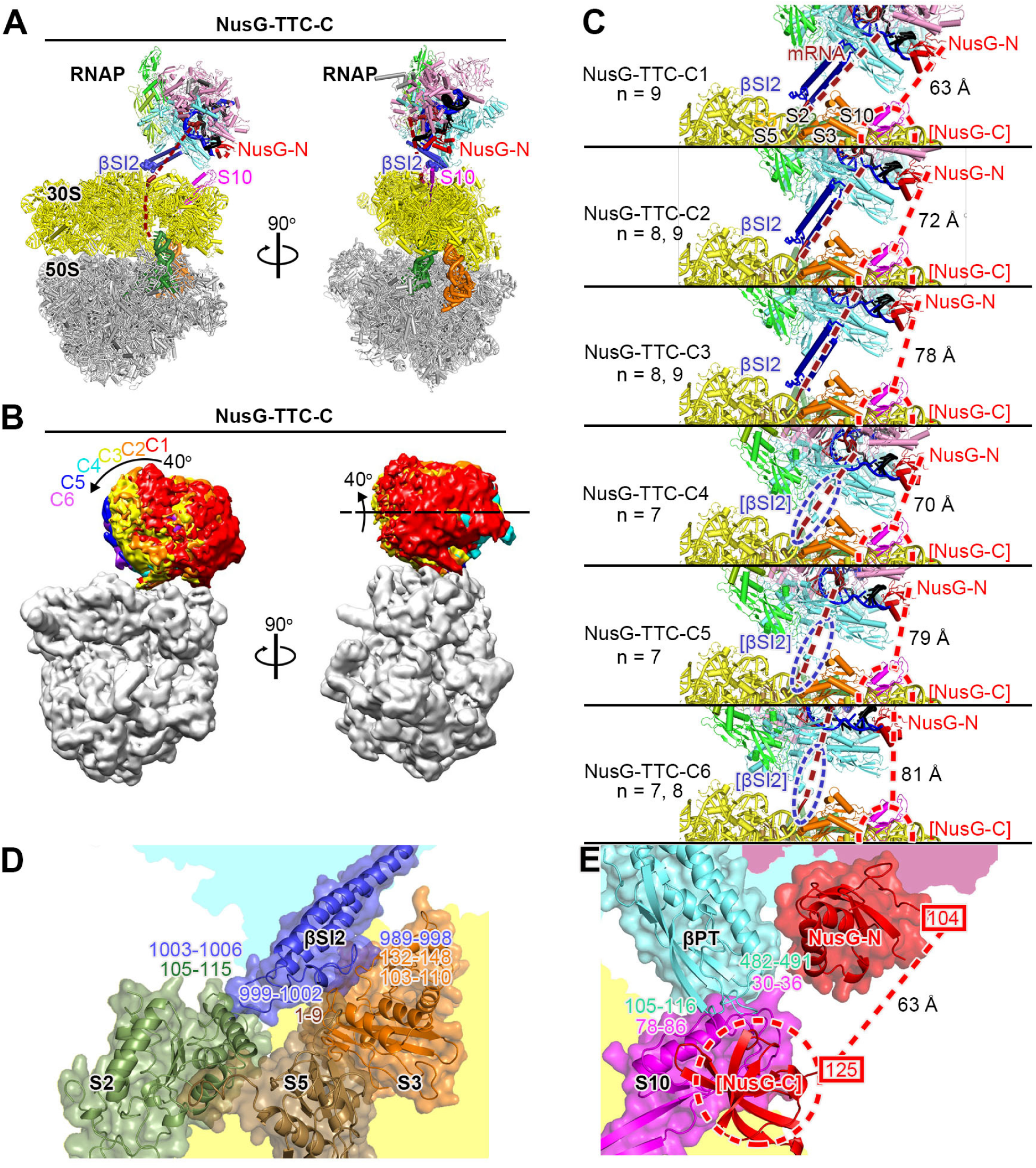
Cryo-EM structure of NusG-TTC-C. **(A)** Structure of NusG-TTC-C (NusG-TTC-C1; 7.6 Å; n = 9; Table S2). Views and colors as in Fig. 2A. **(B)** Subclasses of NusG-TTC-C. Superimposition of NusG-TTC-C subclasses C1-C6 (RNAP EM density colored by subclass; ribosome EM density in gray for subclass C1). View as in (A). **(C)** RNAP-ribosome interactions and potential NusG bridging in NusG-TTC-C subclasses C1-C6. View as in Fig. 3A-left. Ribosomal proteins S2, S3, S5, and S10 in forest green, orange, brown, and magenta. Dark blue ribbons, βSI2; dark blue dashed ovals, disordered βSI2 residues; brick-red dashed line, modelled mRNA; red dashed circle and red dashed line, modelled NusG-C and NusG linker; distances in Å, modelled distances between NusG-N residue 104 and NusG-C residue 125. Other colors as in (A). **(D)** RNAP-ribosome interface in NusG-TTC-C (NusG-TTC-C1; n = 9), showing interaction between βSI2 (dark blue) and ribosomal proteins S2, S3 and S5 (forest green, orange, and brown). Portions of RNAP β and ribosome 30S not involved in interactions are shaded cyan and yellow, respectively. **(E)** Interaction between RNAP β pincer tip (βPT; cyan) and ribosomal protein S10 (magenta) and modelled NusG bridging, with NusG-N interacting with RNAP β’ subunit (pink shading) and RNAP βPT, with NusG-C modelled interacting with S10 (red ribbon and dashed red circle), and modelled linker between NusG-N residue 104 and NusG-C residue 125 as dashed red line. 30S residues not involved in interactions are indicated with yellow shading.

**Fig. S16.**
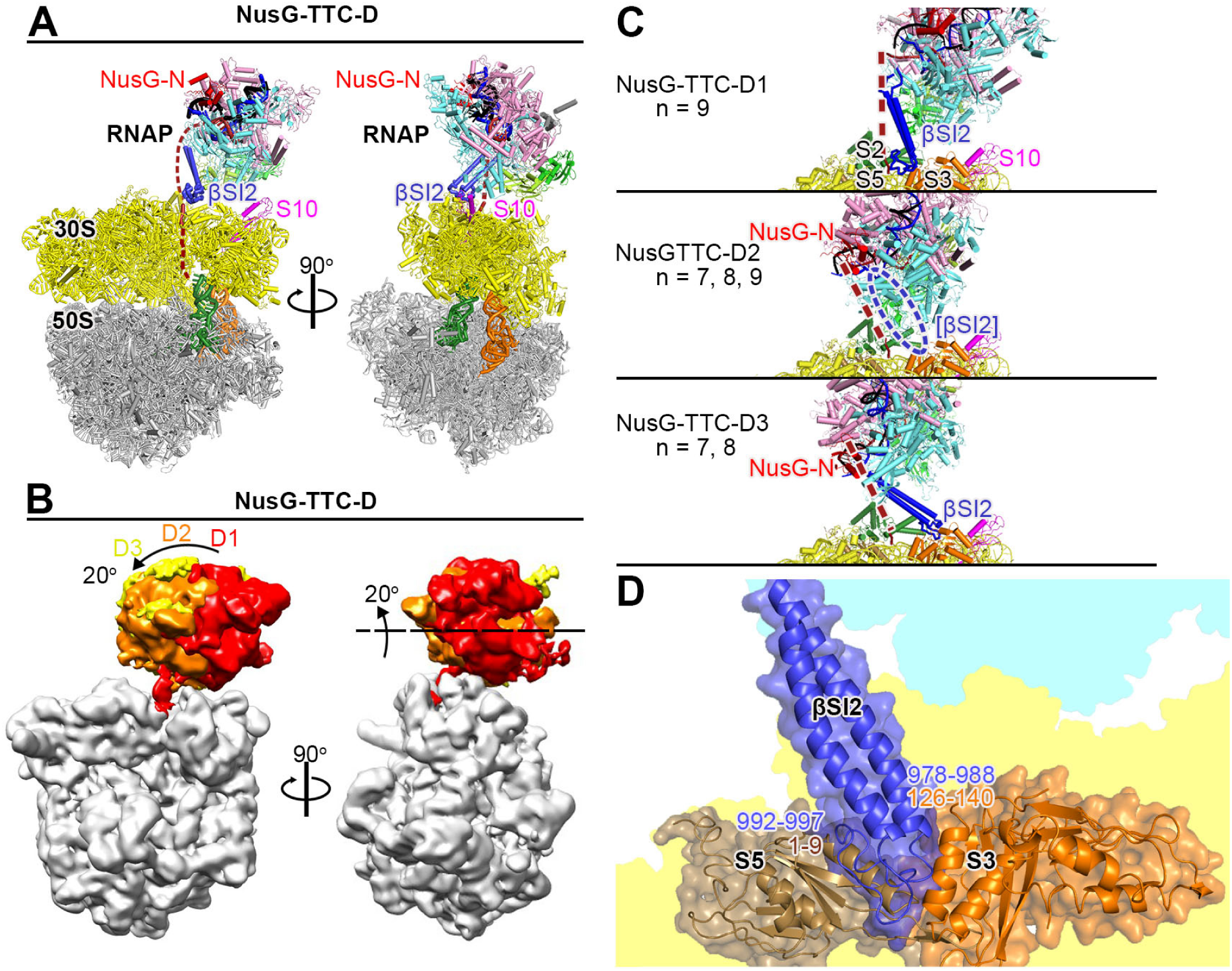
Cryo-EM structure of NusG-TTC-D. **(A)** Structure of NusG-TTC-D (NusG-TTC-D1; 10 Å; n = 9; Table S2). Views and colors as in Fig. 2A. **(B)** Subclasses of NusG-TTC-D. Superimposition of NusG-TTC-D subclasses D1-D3 (RNAP EM density colored by subclass; ribosome EM density in gray for subclass D1). View as in (A). **(C)** RNAP-ribosome interactions in NusG-TTC-D subclasses D1-D3. View as in (A), left. Ribosomal proteins S2, S3, S5, and S10 in forest green, orange, brown, and magenta. Dark blue ribbons, βSI2; dark blue dashed oval, disordered βSI2 residues; brick-red dashed line, modelled mRNA. Other colors as in (A). **(D)** RNAP-ribosome interface in NusG-TTC-D (NusG-TTC-D1; n = 9), showing interaction between βSI2 in dark blue) and ribosomal proteins S2, S3 and S5 (forest green, orange, and brown). Portions of RNAP β subunit and ribosome 30S subunit not involved in interactions are shaded cyan and yellow, respectively.

**Table S1.** Cryo-EM structures: NusG-TTC-A, NusG-TTC-B, and NusA-NusG-TTC-B (n = 4, 5, 6, 7, 8, 9, or 10; with CHAPSO)

| mRNA spacer | TTC class | TTC subclass | cryo-EM facility | particles | resolution | EMDB code | PDB code |
| --- | --- | --- | --- | --- | --- | --- | --- |
| 4 | NusG-TTC-A | TTC-A | NCCAT | 139,302 | 3.7 Å | 21386 | 6VU3 |
| 5 | NusG-TTC-A | TTC-A | Rutgers | 27,378 | 3.7 Å | 21468 | 6VYQ |
| 6 | NusG-TTC-A | TTC-A | Rutgers | 24,582 | 3.8 Å | 21469 | 6VYR |
| 7 | NusG-TTC-A | TTC-A | Rutgers | 29,704 | 3.7 Å | 21470 | 6VYS |
| 8 | NusG-TTC-A | TTC-A | Rutgers | 1,957 | 6.3 Å | 22193 | 6XIJ |
| 5 | TTC-A | TTC-A | Rutgers | 27,650 | 4.1 Å | 21494 | 6VZJ |
| 8 | TTC-A | TTC-A | Rutgers | 10,379 | 3.9 Å |  |  |
| 8 | NusG-TTC-B | TTC-B | Rutgers | 435 | 12.6 Å | 22192 | 6XII |
| 9 | NusG-TTC-B | TTC-B | Rutgers | 6,121 | 4.7 Å | 22142 | 6XDR |
| 10 | NusG-TTC-B | TTC-B | Rutgers | 4,617 | 5.0 Å | 22181 | 6XGF |
| 8 | NusA-NusG-TTC-B | TTC-B1 | NCCAT | 38,958 | 3.2 Å | 22082 | 6X6T |
| 8 | NusA-NusG-TTC-B | TTC-B2 | NCCAT | 45,451 | 3.5 Å | 22084 | 6X7F |
| 8 | NusA-NusG-TTC-B | TTC-B3 | NCCAT | 61,683 | 3.1 Å | 22087 | 6X7K |
| 9 | NusA-NusG-TTC-B | TTC-B1 | Rutgers | 2,558 | 5.9 Å |  |  |
| 9 | NusA-NusG-TTC-B | TTC-B2 | Rutgers | 21,740 | 4.2 Å |  |  |
| 9 | NusA-NusG-TTC-B | TTC-B3 | Rutgers | 11,509 | 4.8 Å | 22107 | 6X9Q |
| 10 | NusA-NusG-TTC-B | TTC-B1 | Rutgers | 4,236 | 4.9 Å |  |  |
| 10 | NusA-NusG-TTC-B | TTC-B3 | Rutgers | 19,968 | 3.7 Å | 22141 | 6XDQ |
| 8 | NusA-TTC-X | TTC-X | Rutgers | 759 | 9.3 Å |  |  |

**Table S2.** Cryo-EM structures: NusG-TTC-A, NusG-TTC-C, and NusG-TTC-D (n = 5, 6, 7, 8, or 9; without CHAPSO)

| mRNA spacer | TTC class | TTC subclass | cryo-EM facility | particles | resolution | EMDB code | PDB code |
| --- | --- | --- | --- | --- | --- | --- | --- |
| 5 | NusG-TTC-A | TTC-A1 | U Michigan | 2,898 | 19 Å |  |  |
| 5 | NusG-TTC-A | TTC-A2 | U Michigan | 6,188 | 14 Å | 21471 | 6VYT |
| 6 | NusG-TTC-A | TTC-A1 | U Michigan | 1,943 | 15 Å |  |  |
| 6 | NusG-TTC-A | TTC-A2 | U Michigan | 9,631 | 13 Å |  |  |
| 7 | NusG-TTC-A | TTC-A2 | U Michigan | 21,801 | 11 Å |  |  |
| 7 | NusG-TTC-C | TTC-C4 | U Michigan | 6,233 | 9.9 Å | 21475 | 6VYX |
| 7 | NusG-TTC-C | TTC-C5 | U Michigan | 6,084 | 9.9 Å | 21476 | 6VYY |
| 7 | NusG-TTC-C | TTC-C6 | U Michigan | 5,913 | 9.9 Å | 21477 | 6VYZ |
| 7 | NusG-TTC-D | TTC-D2 | U Michigan | 6,237 | 8.9 Å |  |  |
| 7 | NusG-TTC-D | TTC-D3 | U Michigan | 6,440 | 8.9 Å | 21485 | 6VZ5 |
| 8 | NusG-TTC-C | TTC-C2 | U Michigan | 7,996 | 7.4 Å |  |  |
| 8 | NusG-TTC-C | TTC-C3 | U Michigan | 4,898 | 9.9 Å |  |  |
| 8 | NusG-TTC-C | TTC-C6 | U Michigan | 6,115 | 9.2 Å |  |  |
| 8 | NusG-TTC-D | TTC-D2 | U Michigan | 5,622 | 9.7 Å |  |  |
| 8 | NusG-TTC-D | TTC-D3 | U Michigan | 2,763 | 12 Å |  |  |
| 9 | NusG-TTC-C | TTC-C1 | U Michigan | 5,979 | 7.6 Å | 21486 | 6VZ7 |
| 9 | NusG-TTC-C | TTC-C2 | U Michigan | 8,928 | 7.0 Å | 21472 | 6VYU |
| 9 | NusG-TTC-D | TTC-C3 | U Michigan | 18,870 | 7.0 Å | 21474 | 6VYW |
| 9 | NusG-TTC-D | TTC-D1 | U Michigan | 4,207 | 10 Å | 21482 | 6VZ2 |
| 9 | NusG-TTC-D | TTC-D2 | U Michigan | 4,960 | 10 Å | 21483 | 6VZ3 |

## Supplementary Movies

**Movie S1. Relationship between NusG-TTC-A and NusG-TTC-B**

Movie shows the ∼70 Å translation and ∼180° rotation that relate NusG-TTC-A to NusG-TTC-B. View and colors as in Fig. 1B (ribosome 50S subunit and tRNAs omitted for clarity).

**Movie S2. Relationship between NusA-NusG-TTC-B subclasses 1, 2, and 3**

Movie shows the up-to-15° rotation on one axis that relate NusG-TTC-B subclasses 1, 2, and 3. Flexible linkage between NusA AR1 and AR2 is shown as light blue circle; flexible linkers between RNAP αCTD^I^ and αNTD^I^ and between RNAP αCTD^I^ and αNTD^I^ are shown as white dashed lines; flexible connectors between RNAP β FTH and rest of RNAP and between RNAP β’ ZBD and rest of RNAP are shown as white circles; flexible linker between NusG-N and NusG-C is shown as red circle. View and colors as in Fig. 1B (ribosome 50S subunit and tRNAs omitted for clarity).

**Movie S3. NusG-TTC-A and 30S-head swivelling**

Movie shows that NusG-TTC-A is incompatible with 30S-head swivelling, the ∼21° rotation of the 30S head (brown) relative to the 30S body (yellow) that occurs during ribosome translocation (*20–22*). View and other colors as in Fig. S11 (ribosome 50S subunit and tRNAs omitted for clarity).

**Movie S4. NusG-TTC-B and 30S-head swivelling**

Movie shows that NusG-TTC-B is compatible with 30S-head swivelling, the ∼21° rotation of the 30S head (brown) relative to the 30S body (yellow) that occurs during ribosome translocation (*20–22*). First clip models RNAP and NusG moving with 30S head; second clip models only RNAP β’ ZBD and NusG-C moving with 30S head. Flexible connectors between RNAP β’ ZBD and rest of RNAP are shown as white circle; f’lexible linker between NusG-N and NusG-C is shown as red circle. View and other colors as in Fig. S11 (ribosome 50S subunit, tRNAs, and mRNA omitted for clarity).

**Movie S5. NusA-NusG-TTC-B subclasses 1, 2, and 3 and 30S-head swivelling**

Movie shows that NusA-NusG-TTC-B is compatible with 30S-head swivelling, the ∼21° rotation of the 30S head (brown) relative to the 30S body (yellow) that occurs during ribosome translocation (*20–22*). First clip for each subclass models RNAP and NusG moving with 30S head; second clip for each subclass models only RNAP β’ ZBD and NusG-C moving with 30S head. Flexible linkers between RNAP αCTD^I^ and αNTD^I^ and between RNAP αCTD^I^ and αNTD^I^ are shown as white dashed lines; flexible connectors between RNAP β FTH and rest of RNAP and between RNAP β’ ZBD and rest of RNAP are shown as white circles; flexible linker between NusG-N and NusG-C is shown as red circle. View and other colors as in Fig. S11 (ribosome 50S subunit, tRNAs, and mRMA omitted for clarity).

**Movie S6. Relationship between NusG-TTC-A subclasses A1 and A2**

Movie shows the ∼60° rotation that relates NusG-TTC-A subclasses A1 and A2. View and colors as in Fig. 1B (ribosome 50S subunit and tRNAs omitted for clarity; mRNA entering 30S subunit shown as in NusG-TTC-A1 for simplicity).

**Movie S7. Relationship between NusG-TTC-A and NusG-TTC-C**

Movie shows the ∼90° rotation on one axis and ∼180° rotation on an orthogonal axis that relate NusG-TTC-A1 to NusG-TTC-C1. View and colors as in Fig. 1B (ribosome 50S subunit and tRNAs omitted for clarity; mRNA entering 30S subunit shown as in NusG-TTC-A1 for simplicity).

**Movie S8. Relationship between NusG-TTC-C subclasses C1, C2, C3, C4, C5, and C6**

Movie shows the up-to-40° rotation that relates NusG-TTC-B subclasses C1, C2, C3, C4, C5, and CB6. View and colors as in Fig. 1B (ribosome 50S subunit and tRNAs omitted for clarity; mRNA entering 30S subunit shown as in NusG-TTC-A1 for simplicity).

**Movie S9. Relationship between NusG-TTC-A and NusG-TTC-D**

Movie shows the ∼45° rotation that relates NusG-TTC-A1 to NusG-TTC-D1. (ribosome 50S subunit and tRNAs omitted for clarity; mRNA entering 30S subunit shown as in NusG-TTC-A1 for simplicity).

**Movie S10. Relationship between NusG-TTC-C and NusG-TTC-D**

Movie shows the ∼180° rotation that relates NusG-TTC-C1 to NusG-TTC-D1. (ribosome 50S subunit and tRNAs omitted for clarity; mRNA entering 30S subunit shown as in NusG-TTC-A1 for simplicity).

**Movie S11. Relationship between NusG-TTC-D subclasses D1, D2, and D3**

Movie shows the up-to-20° rotation that relates NusG-TTC-D subclasses D1, D2, and D3. (ribosome 50S subunit and tRNAs omitted for clarity; mRNA entering 30S subunit shown as in NusG-TTC-A1 for simplicity)

## Notes

### Competing Interest Statement

The authors have declared no competing interest.

